# Massive inversion polymorphisms shape the genomic landscape of deer mice

**DOI:** 10.1101/2022.05.25.493470

**Authors:** Olivia S. Harringmeyer, Hopi E. Hoekstra

**Affiliations:** Department of Organismic & Evolutionary Biology, Department of Molecular & Cellular Biology, Museum of Comparative Zoology and Howard Hughes Medical Institute, Harvard University, 16 Divinity Avenue, Cambridge, Massachusetts 02138, USA

**Keywords:** chromosomal rearrangement, inversions, local adaptation, *Peromyscus*, recombination, structural variation

## Abstract

Chromosomal inversions are an important form of structural variation that can affect recombination, chromosome structure and fitness. However, because inversions can be challenging to detect, the prevalence and hence significance of inversions segregating within species remains largely unknown, especially in natural populations of mammals. Here, by combining population-genomic and long-read sequencing analyses in a single, widespread species of deer mouse (*Peromyscus maniculatus*), we identified 21 polymorphic inversions, which are large (1.5-43.8 Mb) and cause near complete suppression of recombination when heterozygous (0-0.03 cM/Mb). We found that inversion breakpoints frequently occur in centromeric and telomeric regions and are often flanked by long inverted repeats (0.5-50 kb), suggesting that they likely arose via ectopic recombination. By genotyping the inversions in populations across the species’ range, we found that the inversions are often widespread, do not harbor deleterious mutational loads, and many are likely maintained as polymorphisms by divergent selection. Comparisons of forest and prairie ecotypes of deer mice revealed 13 inversions that contribute to differentiation between populations, of which five exhibit significant associations with traits implicated in local adaptation. Together, we found that inversion polymorphisms have a significant impact on recombination, genome structure and genetic diversity in deer mice, and likely facilitate local adaptation across this species’ widespread range.

## Introduction

A longstanding goal in population genetics has been to quantify intraspecific genetic variation, which serves as the substrate for evolutionary change. Since Lewontin and Hubby first characterized protein-sequence variation in *Drosophila pseudoobscura* in 1966, tremendous progress has been made in measuring levels of single nucleotide polymorphisms in a wide diversity of species^1^. However, the prevalence of structural genomic variation, a focus of cytogenetics, still remains largely uncharacterized in the molecular era^2^. Chromosomal inversions, in particular, are an important form of structural variation: inversions can be large (affecting megabases of sequence)^3^ and have been implicated in local adaptation, including differentiation of annual and perennial ecotypes of monkeyflowers^4^, wing-pattern morphs of mimetic butterflies^5^ and mating types of ruffs^6,7^.

Inversions may play a key role in local adaptation because of their effects on recombination. When heterozygous, an inversion will suppress recombination with the non-inverted arrangement, and as a result, can drastically increase linkage disequilibrium between the loci it carries. As such, inversions can act as ‘supergenes’^8^, linking multiple locally adaptive alleles together into co-inherited haplotype blocks, which may be advantageous in the face of gene flow^9–11^. While inversions have been identified across a diversity of species in the context of local adaptation, suggesting that beneficial inversions may be common^3^, few studies have performed unbiased scans across the genome for inversion polymorphisms (but see ^12–15^), raising the question of whether adaptive inversions are the exception or the rule. Thus, characterizing the abundance of inversion polymorphisms – that is, inversions segregating within a species – is a critical step towards quantifying levels of intraspecific genetic variation and understanding how and why inversion polymorphisms establish and are maintained.

Detecting inversion polymorphisms with molecular data has traditionally been challenging (e.g., breakpoints often reside in highly repetitive regions)^16^, but recent advances in long-read sequencing and increased feasibility of population-level genome re-sequencing provide new, powerful approaches for identifying inversions^17,18^. Here, we perform an unbiased genome-wide scan for inversion polymorphisms in the deer mouse, *Peromyscus maniculatus*. The deer mouse is the most abundant and widespread mammal in North America: it has large effective population sizes^19,20^ and a range spanning all major terrestrial habitats, including dense forests and open prairies^21^. Early cytogenetic work suggested that there are visible (large) chromosomal rearrangements segregating in deer mice^22,23^. In this study, we detect 21 large inversion polymorphisms (of which 20 were previously uncharacterized), propose a mechanism by which inversions arise in this species, and survey their distributions both across the species range and across a sharp forest-prairie ecotone. Together, this work reveals proximate and ultimate mechanisms involved in the establishment and maintenance of inversion polymorphisms and suggests a prominent role for these inversions in local adaptation.

## Results

### Identifying inversion polymorphisms

To identify putative inversion polymorphisms, we initially focused on five populations – four deer mouse (*P. maniculatus*) and one oldfield mouse (*P. polionotus*), which is nested within the *P. maniculatus* clade (Figure 1A) – and performed whole-genome re-sequencing (15X coverage with Illumina short-read data) on 15 individuals per population. To identify patterns of genetic variation consistent with inversion polymorphisms, we first characterized local population structure within populations and between population-pairs in 100-kb windows across the genome using local principal component analyses (PCA)^24^ and identified outlier regions (Figure 1B, Figure S1; as described in ^18^). We then focused on genomic regions for which the first principal component separated individuals into three clusters, likely representing the three possible inversion genotypes (Figure 1C, Figure S1) and with the central cluster having highest heterozygosity, consistent with inversion heterozygotes (Figure 1D, Figure S1).

**Figure 1.**
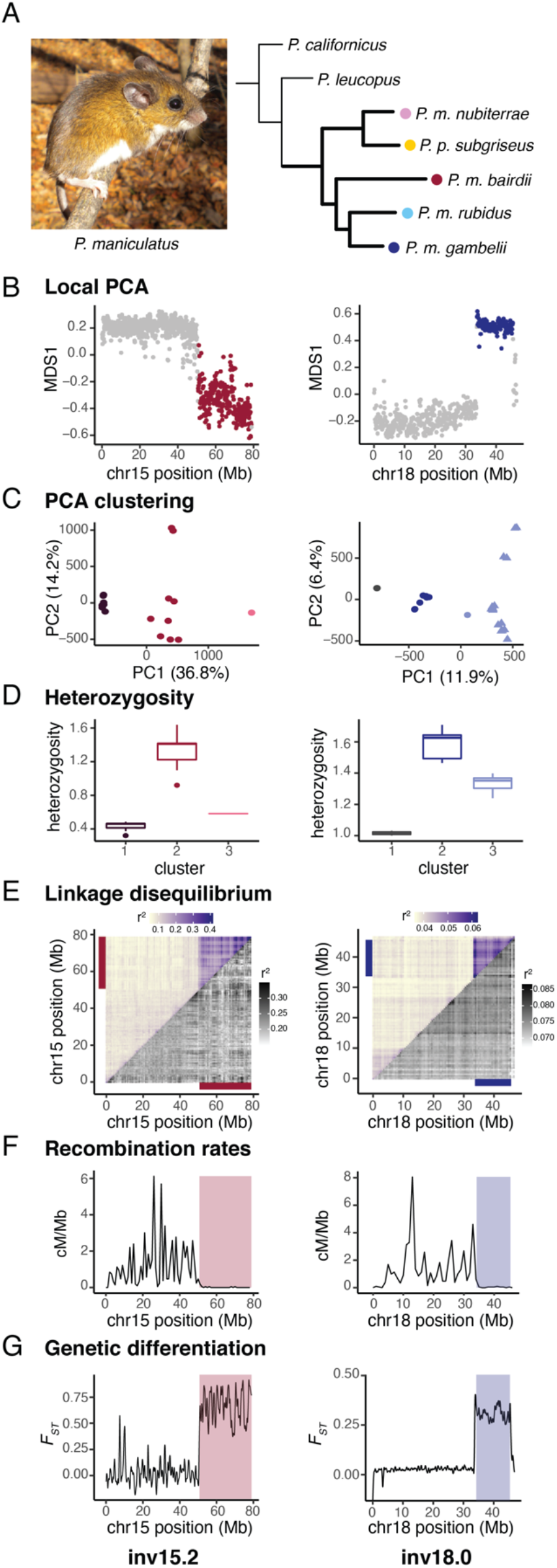
Identifying inversion polymorphisms. (**A**) Left: Photograph of *P. maniculatus* (photo credit E. Kingsley). Right: Phylogenetic tree showing the relationships of five focal populations of *P. maniculatus* (bold branches) and two additional species. Note: *P. polionotus subgriseus* falls within the *maniculatus* clade. (**B-G**) Detection of polymorphic inversions: (**B**) Local PCA for example inversions in *P. m. bairdii* (left) and *P. m. gambelii* (right), where each dot represents a 100-kb window. Distances between local PCA maps are represented by the MDS1 axis, with outlier windows highlighted in color (red or blue). (**C**) Clustering of samples by PCA for entire outlier regions found in (B), assigned using k-means clustering. Right: *P. m. rubidus* shown (triangles) for comparison. (**D**) Heterozygosity of samples by cluster assignments from PCA in (C). Boxplots indicate upper and lower quartiles, with median (center line); whiskers show 1.5x interquartile range; points show outliers. (**E**) Linkage disequilibrium for chromosomes harboring the example inversions, shown as mean *r*^2^ values for paired windows across each chromosome. Mean *r*^2^ values including all samples from PCA clustering (upper triangle) and for only the more common homozygote genotype as determined in PCA clustering (lower triangle). Colored bars highlight outlier regions from (B). Scales for *r*^2^ values provided. (**F**) Recombination rates (cM/Mb) shown for lab-born inversion heterozygotes. Outlier regions found in (B) are highlighted. (**G**) *F*_*ST*_ between homozygous genotypes (clusters 1 and 3 from (C)). Outlier regions found in (B) are highlighted.

To verify that these genomic patterns are driven by suppression of recombination between haplotypes, we next measured linkage disequilibrium (LD) and recombination rates. In wild-caught mice, LD across all genotypes (but not within homozygotes) is elevated within predicted inversion regions (Figure 1E, Figure S1), which suggests that recombination is suppressed between but not within haplotypes. We also estimated recombination rates using lab-raised inversion heterozygotes and found that putative inversion regions showed nearly complete suppression of recombination in heterozygotes (mean recombination per inversion: 0-0.03 cM/Mb; Figure 1F, Figure S1). Together, these results suggest that suppression of recombination is specifically driven by heterozygotes, providing strong evidence that inversion polymorphisms occur in the identified regions. In total, using this approach, we identified 21 inversion polymorphisms, of which 20 are newly discovered, in this species. This is a conservative estimate because our approach was limited to identifying inversions >1 Mb in length with a minimum allele frequency of ∼10%.

Due to their number and size, these inversions alone affect recombination rates on a massive scale. The detected inversions range in size from 1.5 to 43.8 Mb, and in total, span 17.5% of the deer mouse genome. These inversions cause a near complete suppression of recombination in heterozygotes: inversion regions show an average recombination rate of only 0.01 cM/Mb, compared to a genome-wide rate (excluding inversion regions) of 0.80 cM/Mb (Figure S2). We also found no significant correlation between inversion size and recombination rate, highlighting how even the largest inversions almost completely suppress recombination (Figure S2). As a consequence, inversions can trap existing mutations or accumulate new mutations and maintain them in linkage disequilibrium. Indeed, we found that *FST* between inversion and standard haplotypes is elevated in a block-like structure (Figure 1G, Figure S1), suggesting that the inversions partition genetic variation into large haploblocks, shaping patterns of genetic diversity across the deer mouse genome.

### Inversion breakpoints

To localize inversion breakpoints, we performed PacBio long-read sequencing for one individual from each of the five focal populations and created *de novo* genome assemblies at the contig-level (Table S1). By aligning the *de novo* genome assemblies to the deer mouse reference genome (NCBI accession: GCA_003704035.3), we identified breakpoints for 13 out of the 21 inversions (Figure 2A, Figure S3). The eight inversions for which we did not identify breakpoints included five inversions (inv6.0, inv7.0, inv7.1, inv19.0, inv21.0) not represented in homozygotes amongst the PacBio sequenced individuals (Table S2); repetitive sequence likely prevented assembly across breakpoints for the remaining three inversions (inv10.0, inv11.0, inv15.2). Using the *de novo* genome assemblies, we predicted unique centromere locations for 21 out of the 23 autosomes using a 344-bp satellite sequence that localizes to deer mouse centromeres^25^. While centromeres are notoriously difficult to assemble^26^, the *de novo* genome assemblies spanned across multiple predicted centromeres, revealing the highly repetitive nature of centromeric regions, with satellite sequence repeats spanning as much as 1.1 Mb (Figure 2B). Together these data allowed us to precisely map many of the inversions to chromosomes and their positions relative to centromeres.

**Figure 2.**
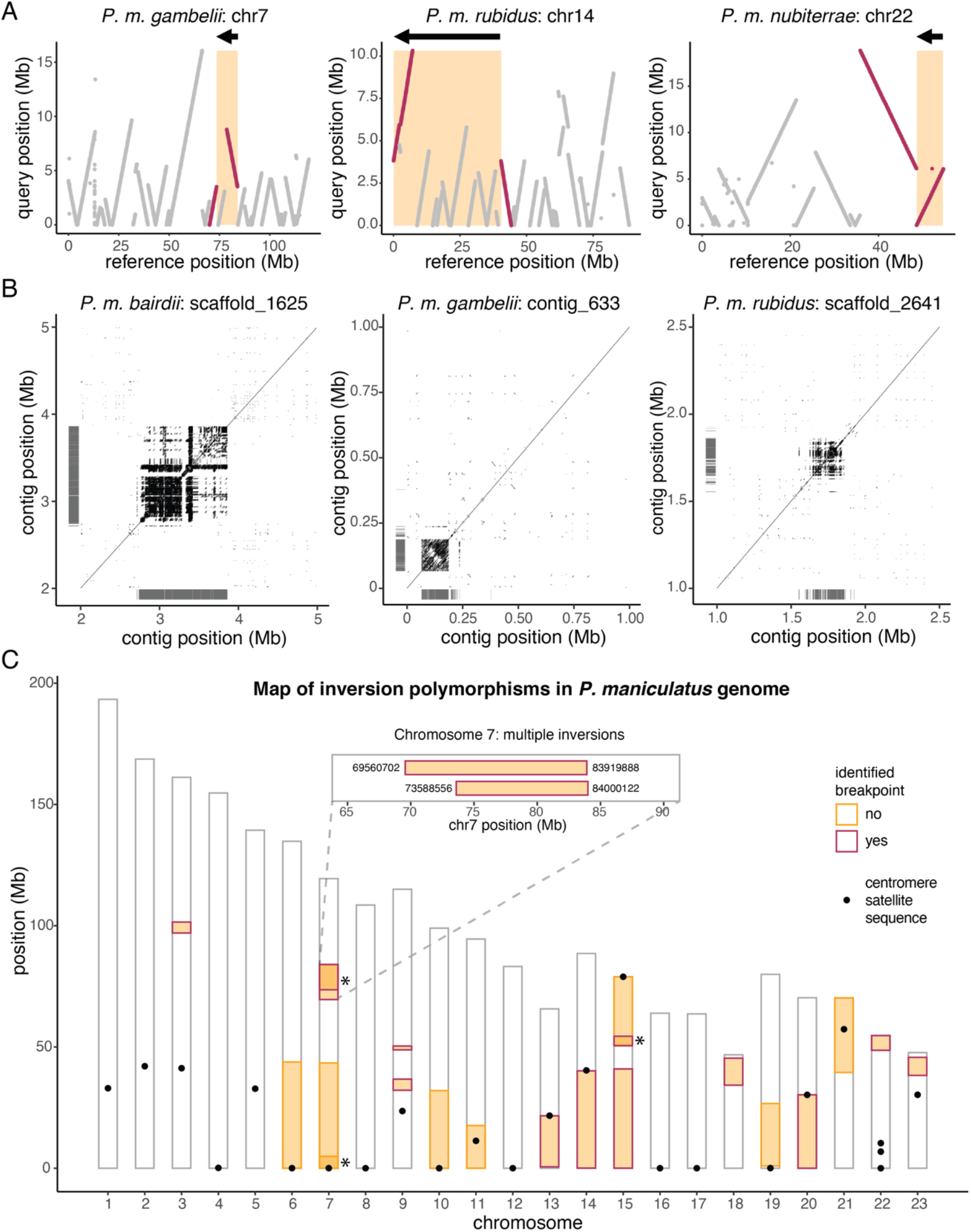
Genome-wide map of inversions. (**A**) Three examples of contigs highlighting inversion breakpoints. Contigs from *de novo* genome assemblies (‘query’, y-axis) were aligned to the *P. maniculatus* reference genome (‘reference’, x-axis) with *nucmer*. Contigs (gray) and those identifying inversion breakpoints (red) are shown. Predicted inversion boundaries are highlighted (orange box), showing predicted inversion (arrow) above. (**B**) Three examples of predicted centromeres in *de novo* genome assemblies. Dotplots show self-v-self alignments with alignment length >100 bp. Location of centromere satellite sequence alignments are shown (gray lines). (**C**) Location of inversions (*n* = 21) across chromosomes, with predicted centromeres (black dots). Asterisks highlight overlapping inversions on chromosomes 7 and 15; inset shows positions of identified breakpoints for two overlapping inversions on chromosome 7.

We found that the distribution of the inversion polymorphisms across the genome is non-random. Of the 21 inversions, 15 are terminal, where the inversion ends within 1.5 Mb of the end of the chromosome (Figure 2C). In addition, nine inversions have breakpoints (predicted or identified) within 1 Mb of the centromere (Figure 2C); since predicted centromeres localize within the three inversions with identified breakpoints (inv13.0, inv14.0, inv20.0) and the other six inversions (inv6.0, inv7.0, inv7.1, inv10.0, inv15.1, inv19.0) are terminal and occur on acrocentric chromosomes, these inversions are likely pericentric (contain the centromere). As such, these nine inversions may be toggling chromosomes between acrocentric and metacentric states, shifting centromere locations by as much as 43 Mb. In addition, these results suggest that, in deer mice, centromeric and telomeric regions are likely to harbor inversion breakpoints.

We also identified multiple genomic regions with recurrent inversion breakpoints. For example, on chromosome 7, we detected two overlapping inversions (inv7.2, inv7.3) with nearly identical breakpoints localizing only 80.2 kb apart (Figure 2C, inset). Using whole-genome alignments between *P. maniculatus* and *P. californicus*, an outgroup, we determined the ancestral versus derived orientation for these two inversions and found that they arose independently rather than as a series of nested inversions. We also identified two inversions on chromosome 15 (inv15.1, inv15.2) with a shared breakpoint (although we localized breakpoints for only one of these inversions with the *de novo* assemblies) and two additional inversions on chromosome 7 (inv7.0, inv7.1) with breakpoints both occurring near the telomere (although we were unable to localize breakpoints for either) (Figure 2C). The recurrence of inversion breakpoints further suggests that certain genomic regions have a greater tendency to participate in the formation of chromosomal rearrangements.

Characterizing the nature of inversion breakpoint regions is critical to understanding how inversions arise and why some genomic regions may be more susceptible to breakpoints. There are two major mechanisms by which inversions form: (1) non-homologous end joining (NHEJ) can create inversions if double stranded breaks occur and the sequence is re-integrated in reverse orientation, and (2) non-allelic homologous recombination (NAHR) can yield inversions if intrachromosomal crossing over occurs between inverted repeats (Figure 3A). For 12 out of the 13 inversions with localized breakpoints, we identified at least one pair of inverted repeats flanking the inversion (Figure 3B). These inverted repeats ranged from 500 bp to 50 kb in length (Figure 3B) and were often duplicated near the breakpoints (Figure 3C, Figure S4). This suggests that the vast majority of inversions for which we identified breakpoints likely arose due to NAHR, as opposed to NHEJ, consistent with the formation of inversions in humans^27^.

**Figure 3.**
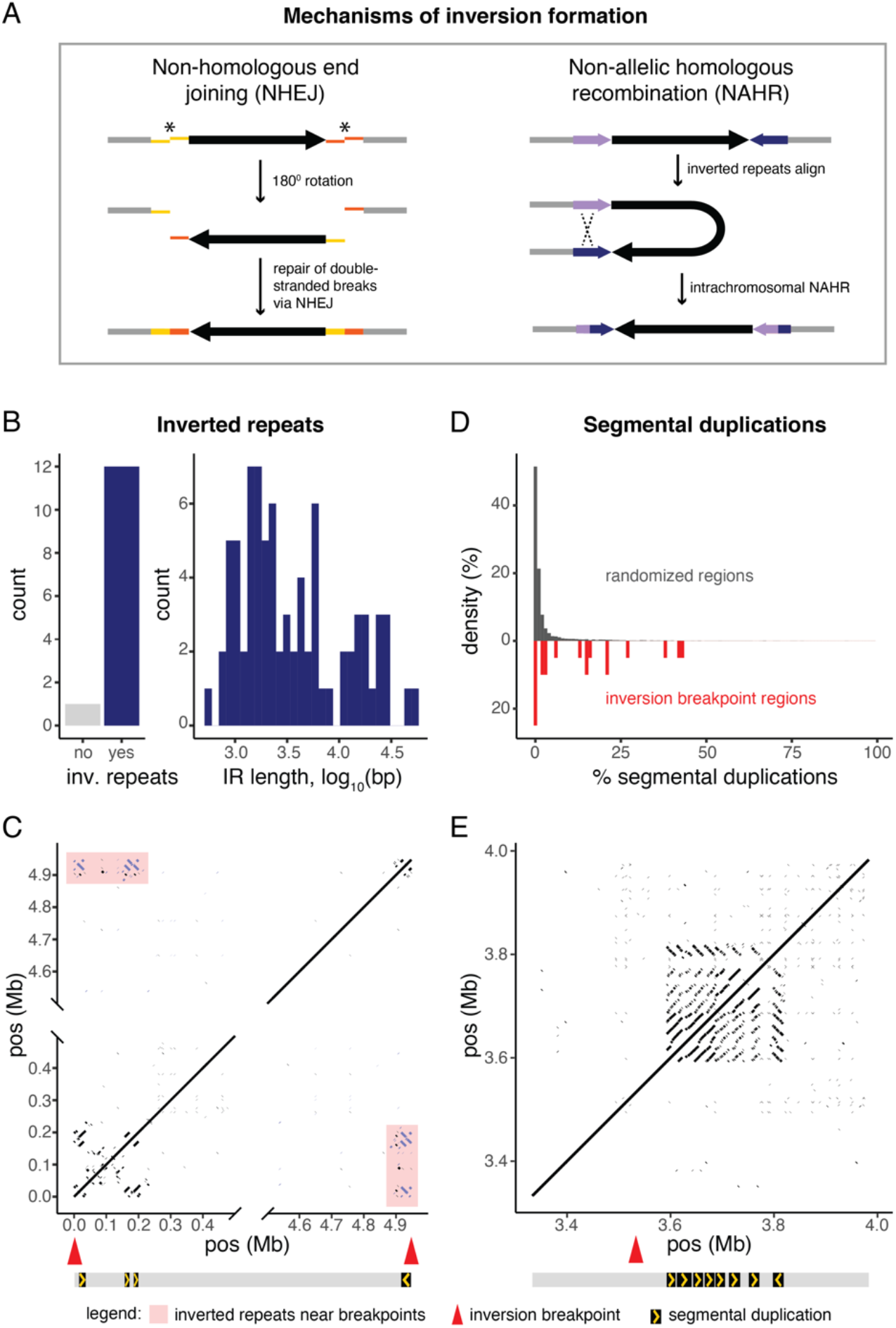
Inversion breakpoints. (**A**) Two primary mechanisms by which chromosomal inversions form. Left: Inversions form when double-stranded breaks (asterisks) occur and sequence is re-incorporated via non-homologous end joining in inverted orientation. Right: Inversions form via non-allelic homologous recombination between inverted repeats (blue and purple arrows) on the same chromosome. (**B**) Number of inversions for which inverted repeats are present or absent within 500 kb of both inversion breakpoints; histogram shows distribution of lengths of identified inverted repeats. (**C**) Example of an inversion on chromosome 9 with inverted repeats. Dotplot shows self-v-self alignments for a long-read assembly contig spanning the inv9.0 inversion. Location of breakpoints (red arrows) shown; only alignments with length >100 bp and within 500 kb of the breakpoints are shown. Inverted repeats mapping to within 500 kb of both breakpoints are shown (purple) and highlighted (pink box). Diagram below shows position and orientation of inverted repeats. (**D**) Percent of sequence assigned as segmental duplications in randomly selected 1-Mb regions across the genome (*n* = 10,000, gray) or within 500 kb of inversion breakpoints (*n* = 20, red). Segmental duplications are defined as regions >1-kb that are duplicated within the selected region, with >70% identity and <70% of sequence masked as common repeats. (**E**) Example of a region harboring segmental duplications near an inversion breakpoint on chromosome 7 (inv7.3). Dotplot shows self-v-self alignments with length >100 bp for a long-read assembly contig spanning the inv7.3 breakpoint; location of breakpoint (red arrow) shown. Diagram below shows position and orientation of a 15-kb region duplicated eight times.

We next explored whether the breakpoints are enriched in repetitive genomic regions. For the 20 localized inversion breakpoints (which excludes six breakpoints at chromosome ends), we used SEDEF^28^ to identify segmental duplications (SDs), defined as duplicated sequence within 500 kb of the breakpoint that is >1 kb in length and contains <70% common repeats. We found that breakpoint regions were significantly enriched for segmental duplications compared to randomized regions genome-wide (KS test: p<0.001); for example, 50% of breakpoints had SD-density in the top 90^th^ percentile of random regions genome-wide (Figure 3D). The repetitive structure of the breakpoints varied, with some breakpoint regions harboring highly structured SDs in tandem (Figure 3E, Figure S4), and other breakpoint regions harboring multiple interspersed SDs (Figure S4). Together, these analyses show that genomic regions with an accumulation of SDs may be prone to chromosomal rearrangements via ectopic recombination in deer mice.

### Frequencies and evolution of inversions

To uncover the evolutionary histories of these inversions, we next characterized their distributions and frequencies across the species range. We first determined the derived inversion arrangement based on genome alignments with an outgroup, *P. californicus*, and then genotyped the inversions in 218 mice from 13 populations (Figure 4A). Most inversions are found in multiple populations: 18 of the 21 inversions were present in at least 3 of the 13 sampled populations (Figure 4B). However, the varying distributions of the inversions suggest that they have differing evolutionary histories (e.g., inversion age, selection): some inversions (e.g., inv14.0) are widespread, whereas others (e.g., inv7.2) are spatially constrained (Figure 4C, Figure S5). Particularly striking is the highly polymorphic nature of many of the inversions (e.g., inv21.0) (Figure 4C, Figure S5), with 16 out of 21 inversions segregating in at least two of the sampled populations (Figure 4B). As such, inversion heterozygotes are common (Figure 4B), indicating that the inversions have a profound impact on recombination rates in the wild.

**Figure 4.**
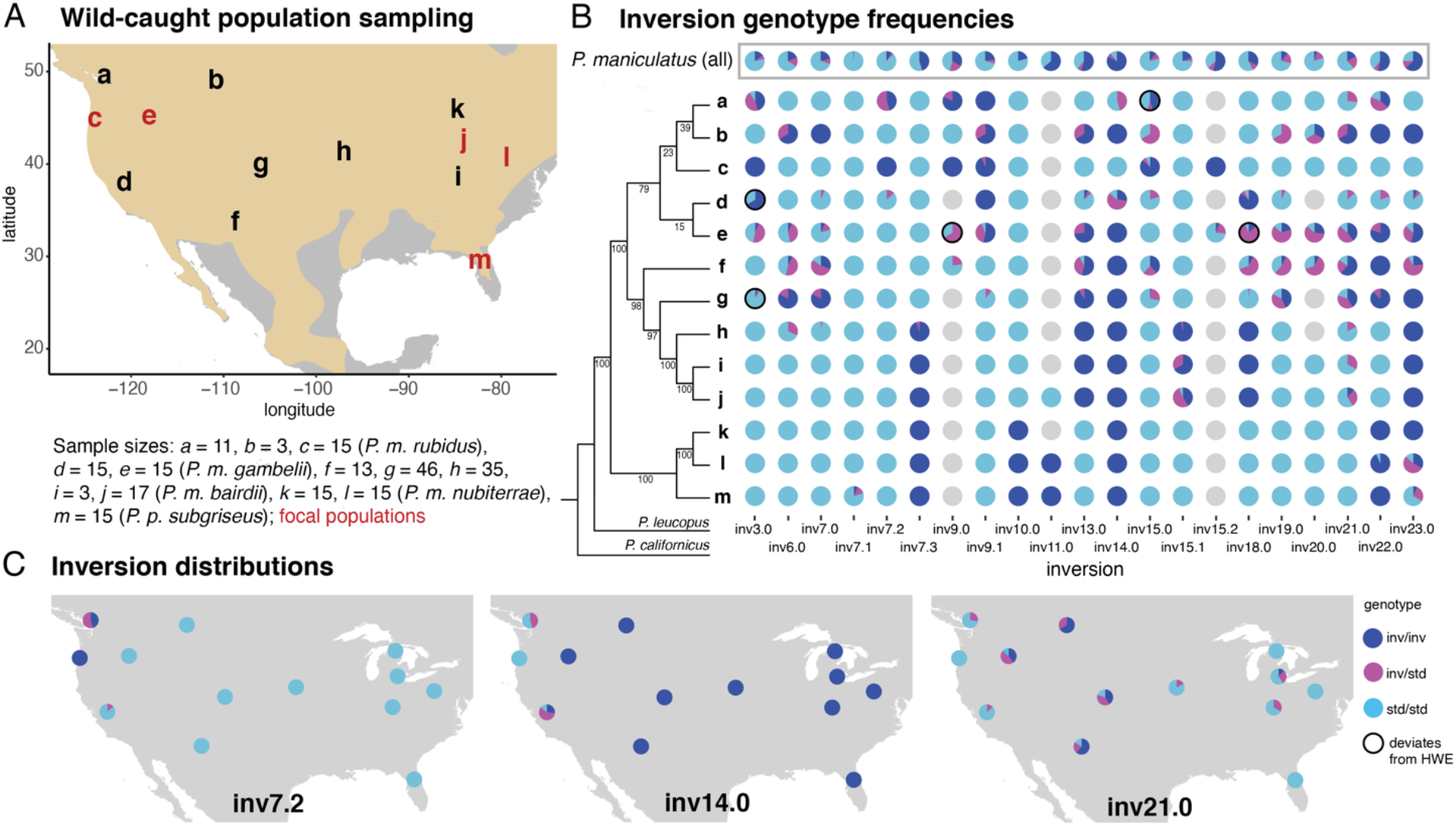
Distributions and frequencies of inversions. (**A**) Locations of all *P. maniculatus* populations sampled (*n* = 13), labelled *a*-*m*; sample sizes given below. The five focal populations are highlighted (red). (**B**) Inversion genotype frequencies for each population: letters correspond to populations in (A). Cladogram (left) shows relatedness (excluding inversion regions) between populations and two outgroups (*P. leucopus, P. californicus*); bootstrap values shown. Inversion genotype frequencies across all *P. maniculatus* populations are shown on top (gray box). Populations that deviate from Hardy-Weinberg equilibrium are shown (*n* = 5, black outline). Missing data shown as gray circles. (**C**) Distributions of three example inversions (inv7.2, inv14.0 and inv21.0) are shown, with inversion genotype frequencies shown for each population. Additional inversion distributions provided in Figure S5.

#### Limited evidence for deleterious effects of inversions

To explore negative consequences of inversions on fitness, we first examined possible deleterious effects on fitness due to inversion breakpoints. If an inversion breakpoint occurs within or near a gene, it may substantially affect the function and/or expression of that gene^29^. We found that significantly fewer inversion breakpoints occurred within protein-coding genes than expected based on the deer mouse gene density (binomial test: *p*=0.004): of the 13 inversions for which we localized breakpoints, only two inversions (inv9.1, inv18.0) had breakpoints occurring within a protein-coding gene (inv9.1 disrupts *1700129C05Rik* intron, inv18.0 disrupts *Slc39a5* coding sequence (left breakpoint) and *Baz2a* intron (right breakpoint)) (Figure 5A). While these two inversions may affect phenotypes through disrupting gene function, the other 11 inversions with localized breakpoints do not disrupt genes (Figure 5A), although their breakpoints may still influence gene expression. Thus, the majority of these 13 inversions are unlikely to confer strongly deleterious effects due to their breakpoints.

**Figure 5.**
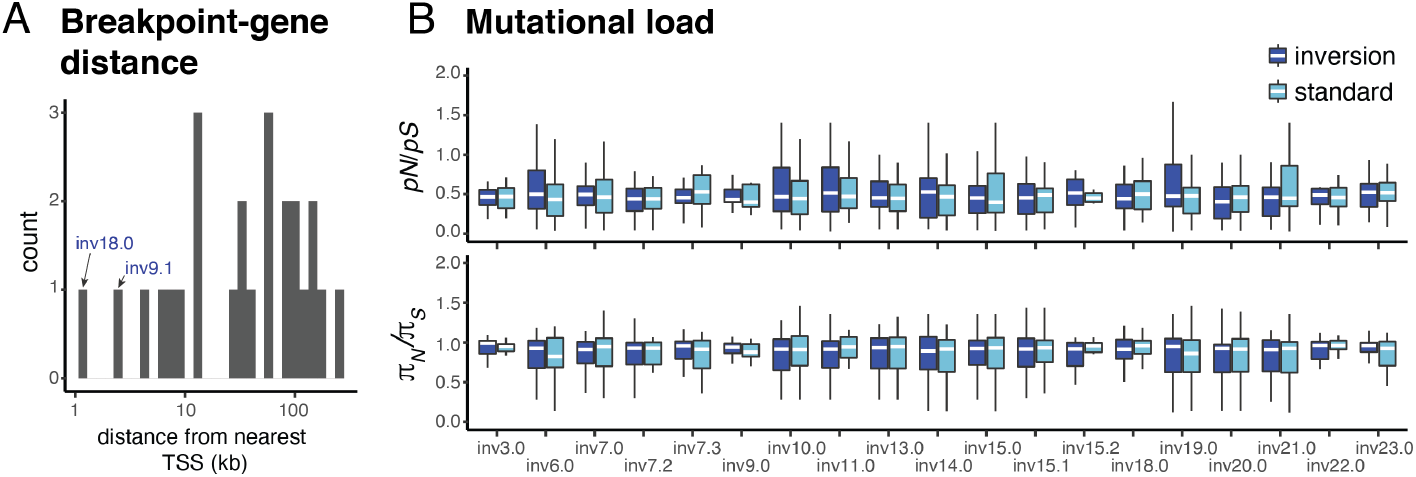
Limited evidence for deleterious effects of inversions. (**A**) Distances between inversion breakpoints (for 13 inversions with localized breakpoints) and the nearest transcriptional start site (TSS). Two inversions (inv18.0, inv9.1) with breakpoints disrupting protein-coding genes are highlighted. (**B**) Ratio of the number of non-synonymous to synonymous polymorphisms (*pN*/*pS*) and nucleotide diversity at non-synonymous versus synonymous sites (*π*_*N*_/*π*_*S*_), computed in 500-kb windows, shown for inversion versus standard haplotypes. Boxplots indicate upper and lower quartiles, with median (center white line); whiskers show 1.5x interquartile range. Differences between inversion and standard haplotypes in *pN*/*pS* and *π*_*N*_/*π*_*S*_ are all non-significant (two-sided t-tests: p>0.05).

We next characterized possible mutational loads carried by the inversions, which may accumulate due to suppressed recombination in inversion heterozygotes^30^. To do so, we tested whether the inversions are enriched for non-synonymous mutations relative to the standard haplotypes. We found that the inversions did not show a significant increase in their proportion of segregating non-synonymous to synonymous mutations (*pN*/*pS*) compared with the standard haplotypes (two-sided t-test: p>0.05 for all inversions) nor did they show a significant increase in nucleotide diversity at non-synonymous versus synonymous sites (*π*_*N*_/*π*_*S*_) compared with the standard haplotypes (two-sided t-test: p>0.05 for all inversions) (Figure 5B). Using non-synonymous mutation accumulation as an estimate of mutational load, these results suggest that the inversions do not harbor a strong deleterious mutational load.

In addition, if inversions accumulate a recessive mutational load, inversion homozygotes should be rare (e.g., in butterflies^31^ and sparrows^32^). In deer mice, however, inversion genotype frequencies are consistent with Hardy-Weinberg equilibrium (HWE): we found only 5 (out of 73) instances in which a segregating inversion significantly deviates from HWE within a population (Figure 4B). This suggests that inversion homozygotes are not strongly underrepresented relative to expectation in populations segregating for a given inversion, which further supports the observation of limited mutational load. We also note that, since most inversion genotype frequencies are consistent with random mating, strong assortative or disassortative mating by inversion genotype is not readily occurring (unlike in the ruff^6,7^ or white-throated sparrow^32^). Together, these lines of evidence suggest that these inversions in deer mice are not associated with strongly negative fitness effects.

#### Multiple inversions contribute to local adaptation

To explore a role of positive selection in the establishment and maintenance of these inversion polymorphisms, we characterized the contribution of inversions to local population differentiation. We took advantage of previous work on two populations, representing forest and prairie deer mouse ecotypes (populations *c* and *e*, Figure 4A), which are well-characterized and widespread ecotypes^19^. Forest and prairie mice show many pronounced phenotypic differences (e.g., coat color, tail length, foot length) despite ongoing gene flow. We previously identified an inversion on chromosome 15 (inv15.0) that contributes to phenotypic divergence between these ecotypes^19^. Returning to this system, we found that multiple, newly identified inversions are also major contributors to differentiation between these populations. Specifically, genome-wide *FST* is low between ecotypes (genome-wide forest-prairie *FST*: 0.03±0.03) due to high migration rates^19^, yet we found multiple ‘genomic islands of divergence’ that show remarkable overlap with identified inversion polymorphisms (inversion regions forest-prairie *FST*: 0.26±0.16) (Figure 6A). For 13 inversions, the ecotypes differ by >50% in their inversion frequencies. Using forward-genetic simulations in SLiM^33^, we found that for a locus to be maintained at >50% frequency difference between the forest and prairie ecotypes given high gene flow, it is most likely evolving under divergent selection (Figure S6), implicating these 13 inversions in local adaptation.

**Figure 6.**
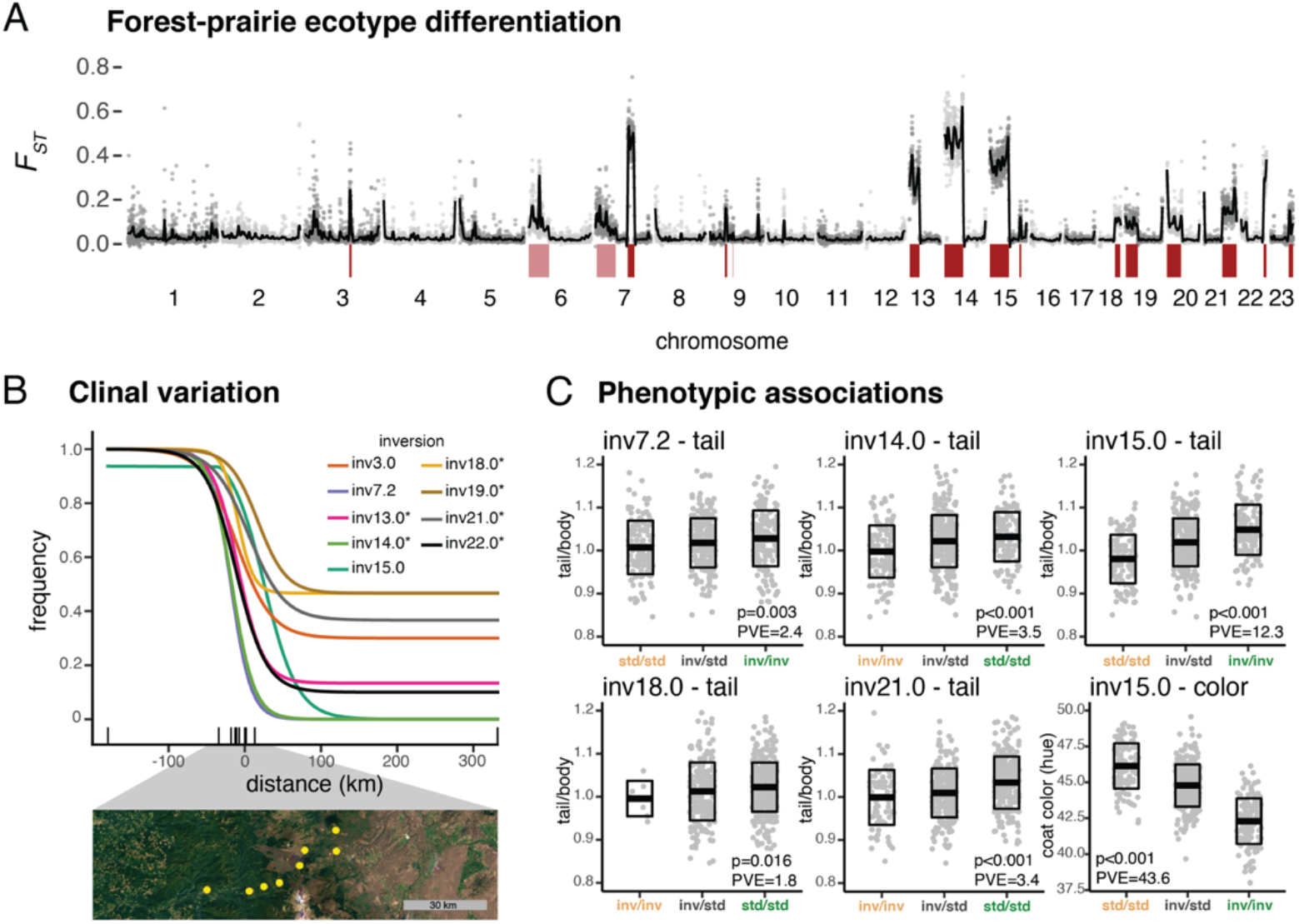
Inversions involved in local adaptation. (**A**) *F*_*ST*_ between forest and prairie ecotypes found in western North America. Gray dots show *F*_*ST*_ from 100-kb windows, with smoothed *F*_*ST*_ as black lines. Inversions with allele frequency difference >50% between forest and prairie ecotypes are shown in dark red; inversions with allele frequency difference >10% in pink. (**B**) Inversion or standard arrangement frequencies across an environmental gradient found between the forest and prairie populations, with clines fit using *hzar*. Sampled populations are shown with black ticks along x-axis, with forest population as left-most tick and prairie population as right-most tick. Clines show frequency of the forest allele: asterisk indicates standard arrangement is more common in forest population, no asterisk indicates inversion arrangement is more common in forest population. Satellite image across the environmental gradient is shown below (GoogleMaps), with sampled sites in yellow. (**C**) Inversions with significant associations with tail length or coat color in a forest-prairie F_2_ intercross. F_2_ hybrids are grouped by inversion genotype, with genotype colored by whether it is more common in the prairie (tan) or forest (green) population. Boxes show mean±sd. P-values (Bonferroni corrected) and percent variance explained (PVE) from linear models for each association are shown. Tail/body = ratio of tail length to body length; coat color is measured by hue, shown in degrees.

The distributions of these inversions across a forest-prairie habitat gradient further support their role in adaptation. Specifically, we genotyped the 13 polymorphic inversions in 136 samples across an environmental gradient and found that nine inversions show steep changes in frequency across the forest-prairie habitat transition (Figure 6B, Figure S7), suggesting that these inversions may be favored in alternate habitats. Furthermore, five inversions (inv7.2, inv14.0, inv15.0, inv18.0, inv21.0) are significantly associated with an ecotype-defining trait, tail length, in lab-raised F2 hybrids^19^, and for all five, the forest arrangement is associated with longer tails (Figure 6C), consistent with long tails being important for balance in arboreal habitats^34^. Inv15.0 was also previously found to be significantly associated with coat color, a second ecotype-defining trait^19^ (Figure 6C). Together, these results suggest that inversions may be a key source of genetic variation differentiating locally adapted deer mouse populations, with divergent selection likely playing a role in maintaining the inversions as polymorphisms within this species.

## Discussion

Technological advances in genome sequencing have recently opened new opportunities for characterizing intraspecific structural variation. For example, the ability to perform population-level whole-genome re-sequencing allows for signatures of large structural variants such as chromosomal inversions to be more easily detected^18^. This approach recently has been successful in identifying inversions in sunflowers^15,35^, seaweed flies^14^, and now in deer mice. In addition, long-read sequencing has also greatly facilitated the detection and classification of structural variants. For example, here we found that the inversion breakpoints reside in highly repetitive genomic regions, harboring an enrichment of segmental duplications, similar to other mammalian species (i.e., humans and great apes^16^). The repetitive nature of mammalian inversion breakpoints likely explains why breakpoints are so challenging to detect with short-read sequencing data alone as well as with long-read data if read length or coverage is insufficient to resolve repeat regions, as we suspect is the case for the deer mouse inversions for which we failed to localize breakpoints. Future work, combining these two approaches – to perform population-level long-read genome sequencing – will further our ability to detect structural variation within a diversity of species^17^.

In discovering deer mouse inversion polymorphisms, we find they have an interesting distribution in the genome: a majority of the inversions occur terminally, most of which involve breakpoints near centromeres. The inversions with breakpoints adjacent to centromeres are likely shifting centromere locations from the middle of the chromosome to the end of the chromosome (and vice versa), transforming chromosomes between metacentric and acrocentric states. This result can explain the longstanding observation that deer mice vary in number of acrocentric chromosomes^22,23^. Despite this large variation in chromosome structure, deer mice (and more generally, the *Peromyscus* genus) have a strongly conserved chromosome number (diploid *n* = 48)^23^. Unlike in other rodents such as the house mouse that harbors Robertsonian fusions^36^, the large rearrangements involving centromeres occur primarily within, and not between, chromosomes in deer mice.

One hypothesis for why deer mouse inversions tend to involve telomeric and centromeric regions is that inversion breakpoints arise more frequently in these regions: genomic regions near centromeres and telomeres can harbor an excess of segmental duplications (as well as other repeats), which may facilitate ectopic recombination^37^. A second hypothesis is that inversions with breakpoints in telomeric or centromeric regions are less likely to be removed by purifying selection than inversions that occur in other genomic regions: breakpoints that occur near centromeres and telomeres may be unlikely to have deleterious effects since these regions tend to be gene-sparse^37^. Indeed, none of the inversion breakpoints we found near centromeres (and only one near a telomere) disrupt protein-coding sequences. Terminal inversions may also be less likely than non-terminal inversions to have strong underdominant effects, which often occur due to inversion loops that form in heterozygotes during meiosis^3^. If an inversion lacks homologous sequence on one side, such as in a terminal inversion, loop formation may be prevented. Previous evidence from deer mice suggests that inversion loop formation is rare in putative terminal inversions^38^. Thus, deer mouse inversions involving telomeres and centromeres may confer fewer deleterious costs associated with breakpoint effects and underdominance than inversions occurring in the rest of the genome.

Inversions are a particularly interesting form of structural variation because of their effects on recombination. Consistent with inversions in other organisms^3^, inversions in deer mice, when heterozygous, suppress recombination across their entire lengths. Therefore, the number and sizes of the inversions is remarkable: 21 detected inversion polymorphisms, with mean length 20.0 Mb, affect a total of 420 million DNA basepairs in the deer mouse genome. Due to the widespread distributions of most of these inversions and the common occurrence of inversion heterozygotes we detected in natural populations, inversions substantially shape the recombination landscape of deer mice.

Recombination plays an important role in evolution, through creating new combinations of alleles and increasing the efficiency of natural selection^39^. In particular, through uncoupling deleterious and beneficial mutations, recombination reduces Hill-Robertson interference and facilitates the elimination of deleterious mutations and the spread of beneficial mutations^40,41^. Given the benefits of recombination, the abundance of inversions presents a paradox. With reduced efficacy of purifying selection in the absence of recombination, the expectation is that inversions will accumulate a deleterious mutational load (when inversion heterozygotes are common)^30^, which will limit their spread^42^. In deer mouse inversions, however, we did not find evidence for the accumulation of mutational load, consistent with a recent study in sunflowers^43^. In both deer mice and sunflowers, inversion homozygotes are common^43^; since recombination proceeds uninterrupted in inversion homozygotes, deleterious mutations can efficiently be removed once an inversion reaches substantial allele frequency^30^, especially if effective population sizes are high, as in many populations of deer mice (e.g., *Ne* ≈ 4e6 in a single population^19^). We hypothesize that these inversions largely evaded mutational load accumulation by quickly spreading in deer mice, whose large population sizes could facilitate effective purifying selection in inversion homozygotes, noting that gene conversion between inversion and standard haplotypes may also play a role in reducing deleterious mutational load^30^.

A major hypothesis for the maintenance of inversion polymorphisms is the ‘local adaptation hypothesis’ which posits that, when a population is locally adapting in the face of gene flow, suppressed recombination between multiple beneficial mutations can be advantageous, reducing the strength of selection necessary to establish and/or maintain each mutation in migration-selection equilibrium^9–11^. Since deer mice are found continuously across a wide range of habitats, they are subjected to a range of selective pressures, likely with ongoing gene flow. Our results support an important role for divergent selection in maintaining inversions as polymorphisms within the species at large. In particular, we found that 13 inversions, including one previously identified^19^, are segregating between forest and prairie deer mouse ecotypes with high allele frequency differences and are likely subject to habitat-associated divergent selection, which is consistent with multiple inversions differentiating ecotypes in a diversity of species such as snails^44^, cod^45^, sunflowers^35^, and sticklebacks^46^. While it remains an open question whether the inversions segregating between these forest-prairie ecotypes are advantageous because of their suppression of recombination, the high levels of migration between the forest and prairie populations suggest that increased LD between adaptive alleles may be particularly beneficial in this system^19^. In addition, five of these inversions have significant effects on tail length, and thus variation in this ecotype-specific trait is largely partitioned into inversions, which is consistent with the evolution of concentrated genetic architectures in the face of gene flow^47^.

A concrete understanding of the prevalence and significance of inversion polymorphisms specifically, and structural variation more generally, remains largely elusive across natural populations of organisms, especially mammals^48^. We find that inversion polymorphisms are remarkably abundant in deer mice. Whether the abundance of inversion polymorphisms in deer mice is unique or representative of mammalian species will require similar investigations across additional species. Nevertheless, this work highlights the critical role of inversions in shaping patterns of recombination, genetic diversity and chromosomal structure in the deer mouse and suggests that inversions may play an even more important role in local adaptation than previously appreciated.

## Methods

### Population sampling and sequencing

#### Focal population sampling

We focused our initial analyses on five populations of *P. maniculatus*, each representing a distinct subspecies (*P. m. rubidus, P. m. gambelii, P. m. bairdii, P. m. nubiterrae, P. p. subgriseus*). Tissues for 15-17 wild-caught mice per population were collected in Siuslaw National Forest, Oregon, USA (*P. m. rubidus*)^19^, Baker City, Oregon, USA (*P. m. gambelii*)^19^, Derry, Pennsylvania, USA (*P. m. nubiterrae*)^49^, Ocala National Forest, Florida, USA (*P. p. subgriseus*)^50^ and Bridgewater, Michigan, USA (*P. m. bairdii*; obtained from the University of Michigan). All samples used in this study are listed in Table S3.

#### Whole-genome re-sequencing and variant calling

To generate whole genome re-sequencing data, we first extracted DNA from ∼20mg of liver tissue and generated sequencing libraries using Illumina DNA library preparation kits. We sequenced the resulting libraries using 150 bp paired-end sequencing on an Illumina NovaSeq S4 flowcell. Following demultiplexing, we mapped sequencing reads to the *P. maniculatus bairdii* reference genome (NCBI accession: GCA_003704035.3) using BWA-MEM. For three populations, we accessed published re-sequencing data: *P. m. rubidus* and *P. m. gambelii*^19^ (NCBI: PRJNA688305) and *P. p. subgriseus*^50^. To call variant sites, we used *HaplotypeCaller* (GATK3.8) on each sample with the default heterozygosity prior (-hets = 0.001) and –ERC GVCF to produce per sample gVCFs. Then, we ran *GenotypeGVCFs* (GATK3.8) to jointly genotype the samples. We performed hard filtering of SNPs based on GATK best practices (filtering variants with QD <2.0, FS >60.0, MQ <40.0, MQRankSum < -12.5, ReadPosRankSum < -8.0) using *VariantFiltration*.

### Identifying inversions

#### Local PCA

To identify genomic regions with outlier population structure, we performed local principal component analyses (PCA) with the *lostruct* package^24^ in R on each of the five focal populations and for all focal-population pairs. Using *lostruct*, we performed local PCA for 100-kb windows with step-size of 100 kb. We then computed the distance between PCA maps (with the top 2 PCs) using *pc_dist* function with default parameters and visualized these distances using multi-dimensional scaling (MDS) with the *cmdscale* function with two MDS axes.

To identify genomic regions with unusual population structure, we scanned for consecutive 100-kb windows which showed similar population structure to each other and distinct population structure from the rest of the chromosome. To do so, we first performed k-means clustering of the 100-kb windows in the MDS space, defined by MDS1 and MDS2 axes, using number of clusters from *k*=2 to *k*=10. To determine the best *k*, we used the silhouette score, which is an averaged measure of the dissimilarity between an observation and its neighboring cluster. We chose the *k* with maximum silhouette score, and assigned 100-kb windows to the cluster determined by the k-means clustering for the chosen *k*. We next calculated the z-score for the MDS1 score for each 100-kb window and selected genomic regions with consecutive windows belonging to the same cluster, in which at least 10 consecutive windows had z-score >1.5.

#### PCA and heterozygosity

For each identified outlier region, we performed PCA on the entire region using *scikit-allel* v1.3.2 (https://github.com/cggh/scikit-allel). For *scikit-allel* analyses, we created zarr objects from the whole-genome re-sequenced vcfs using *allel*.*vcf_to_zarr*. We then performed PCA using all SNPs in the region, with the function *allel*.*pca*, with n_components=10, scaler=‘patterson’, and ploidy=2. K-means clustering of samples in PC1 v. PC2 space was performed in R with *kmeans*, following the approach detailed in Todesco et al.^15^, where samples were assigned to three clusters, setting the cluster starting positions as the minimum, maximum and middle value for PC1 scores to prevent clustering from being influenced by unequal number of samples per cluster. When clustering into three groups failed, we tried clustering into two groups, which would be the case if only two inversion genotypes are present. In a few cases (*n* = 4), we manually re-assigned clusters for samples when k-means clustering had clear misassignments. For each outlier region identified, we also computed heterozygosity (reported as percent of sites that are heterozygous) for every sample in the relevant populations, using *count_het* in *scikit-allel*. Finally, we selected putative inversions to be outlier regions for which samples clustered into three distinct groups along PC1 with high heterozygosity for the middle cluster. We also included an additional four regions for which samples clustered into only two distinct groups along PC1, but signatures of recombination suggested the presence of an inversion (see below).

#### Linkage disequilibrium

For each putative inversion, we computed linkage disequilibrium (LD) across the chromosome harboring that putative inversion using either (1) all samples belonging to the population or population pair from which the putative inversion was identified or (2) only the samples homozygous for the more common haplotype, based on the PCA clustering. To compute LD, we first used bcftools to subset the vcf by sample and chromosome. We then used vcftools to filter for SNPs with MAF >5% (--maf 0.05) and number of missing genotypes = 0 (--max-missing-count 0), and thinned SNPs to at most 1 SNP per 1 kb (--thin 1000). We computed linkage disequilibrium with vcftools geno-r2. Finally, we used the script emerald2windowldcounts.pl (https://github.com/owensgl/reformat, https://github.com/owensgl/haploblocks) to calculate the mean *r*^2^ between 500-kb windows (i.e., for a given set of two 500-kb windows, the mean *r*^2^ across all pairwise SNP comparisons between the two windows was computed).

#### Recombination rates

We estimated recombination maps for both the whole-genome and within-inversion regions, using lab-raised F2 hybrids from previous intercrosses between two population pairs: *P. m. rubidus* x *P. m. gambelli*^19^ and *P. m. bairdii* x *P. p. subgriseus*^51^, which yielded a total of 547 and 1061 F2 hybrids, respectively. Using ddRAD-sequencing data of F2 hybrids, we determined ancestry and the location of recombination breakpoints in the F2 hybrids using the multiplexed genotyping pipeline (see ^19^ for details). For the *P. m. rubidus* x *P. m. gambelli* intercross, we genotyped the founders (*n* = 4) and F1 hybrids (*n* = 49) of the intercross for the inversions (see section ‘Genotyping samples for inversions’) to ensure that only F2 hybrids that were offspring of F1 inversion heterozygotes were used for computing recombination rates within inversion regions. All inversions analyzed in the *P. m. bairdii* x *P. p. subgriseus* intercross were fixed between the founders. Five inversions (inv7.0, inv7.3, inv9.1, inv15.2, inv20.0) were not represented by heterozygous F1 hybrids, and so we are unable to characterize recombination rates for these inversions.

#### Genetic differentiation

To measure genetic differentiation between inversion and standard haplotypes across each identified inversion, we computed *FST* between predicted homozygote genotypes (clusters 1 & 3 from PCA clustering) using *scikit-allel*. We performed sliding window *FST* analyses for 10-kb windows with 10 kb step size using *scikit-allel* with the *windowed_hudson_fst* function and visualized *FST* with *loess* smoothing in R.

To analyze genome-wide genetic differentiation between forest (*P. m. rubidus*) and prairie (*P. m. gambelii*) ecotypes, we computed *FST* between forest and prairie populations in 100-kb windows across the genome with step size of 100 kb, using *scikit-allel* with the *windowed_hudson_fst* function.

### Localizing inversion breakpoints

#### PacBio long-read sequencing and de novo genome assembly

We performed long-read sequencing on five individuals, one from each focal population. For long-read sequencing, we used lab-colony raised mice. First, we extracted high-molecular weight (HMW) DNA from 200 uL fresh blood using the MagAttract HMW DNA mini kit (Qiagen), following the Whole Blood protocol. We quantified the resulting DNA using a Genomic DNA ScreenTape on the Tapestation 4200 (Agilent). Library preparations and sequencing were performed at the University of Washington’s PacBio Sequencing Core. In brief, libraries were prepared with the SMRTbell Express Template Prep Kit 2.0 (PacBio). We performed a size selection of 30 kb for the *P. m. rubidus, P. m. nubiterrae* and *P. m. bairdii* samples using the BluePippin (Sage Science); we did not perform any size selection for the *P. m. gambelii* and *P. p. subgriseus* samples since total library mass was below 500 ng. We then sequenced each on a Sequel II SMRTcell 8M (PacBio), the *P. m. rubidus P. m. nubiterrae* and *P. m. bairdii* samples with a 15-hour movie and the *P. m. gambelii* and *P. p. subgriseus* samples with a 30-hour movie.

We converted the bam files from each movie to fastq files using bam2fastx (PacBio). We then used *flye*^52^ to create *de novo* genome assemblies at the contig-level for each population. The *flye* assembler uses a repeat graph to assemble across repetitive genomic regions, a critical feature for localizing inversion breakpoints which often occur in repetitive genomic regions. To reduce run time, we down-sampled to 40X coverage (-asm-coverage=40) for initial disjointing assembly but otherwise ran the assembler with default parameters. Genome qualities are reported in Table S1.

To genotype each PacBio sample for the inversions, we first mapped the PacBio fastq files to the *P. maniculatus* reference genome using *ngmlr*^53^. Then, we used *longshot*^54^, a long-read-specific variant caller, to call variants for each sample. We merged the variant calls with the whole-genome re-sequencing vcfs and performed PCA for each inversion region, which allowed us to genotype the PacBio samples for the inversions (reported in Table S2) (for details, see section ‘Genotyping samples for inversions’).

#### Inversion breakpoint identification

We aligned the PacBio genome assemblies to the *P. maniculatus bairdii* reference genome using *nucmer* (*mummer*)^55^ with default parameters. Because of possible reference genome errors, we re-oriented any scaffolds in the reference genome that were misoriented relative to the *P. m. bairdii* long-read assembly (i.e., we identified signatures of inversions or translocations in the *P*.*m. bairdii* long-read assembly relative to the reference genome and resolved these regions to match the *P. m. bairdii* long-read assembly). Thus, all inversion analyses were relative to the *P. m. bairdii* long-read assembly. We also aligned published *P. californicus*^56^ (NCBI accession: GCA_007827085.2) and *P. leucopus*^57^ (NCBI accession: GCA_004664715.2) genomes as well as previously assembled *de novo* genomes for *P. m. rubidus* and *P. m. gambelii*^19^ from *canu* (a complementary long-read genome assembler to *flye*) to the *P. maniculatus* reference genome using *nucmer*.

For each inversion, we scanned for evidence of inversion breakpoints. To do so, we filtered for *nucmer* alignments >4 kb in length (or >10 kb for *P. californicus, P. leucopus* alignments). Inversion breakpoints are identifiable if (1) a contig spans across the inversion region and maps to the reference genome in opposite orientation within the inversion region, or (2) a contig spans only part of the inversion region and maps to the reference genome in opposite orientation to the flanking region of the other end of the inversion. We thus identified contigs that showed signatures of inversions in predicted inversion regions and identified breakpoint positions based on the PacBio assembly alignments to the *P. maniculatus* reference genome. In addition, we identified breakpoints for one of the predicted inversions based on the *P. leucopus* genome alignment to the *P. maniculatus* reference genome, and one of the predicted inversions based on the *P. californicus* genome alignment to the *P. maniculatus* reference genome.

#### Determining derived arrangement

For each inversion polymorphism, we determined which arrangement was ancestral (standard) versus derived (inversion) based on the whole-genome alignments between *P. californicus* (outgroup) and *P. maniculatus*. We evaluated whether the *P. californicus* reference genome was inverted relative to the *P. maniculatus* reference genome for each inversion region, and we assigned the *P. californicus* orientation to be the ancestral, standard arrangement.

#### Predicting centromere locations

*Peromyscus* are known to have satellite sequences that map to centromeres, specifically a 344-bp satellite sequence (NCBI accession: KX555281.1) localizes to *P. maniculatus* centromeres^25^. We used *blastn* (blast v2.2.29) to map this satellite sequence to the *P. maniculatus* reference genome and to each PacBio genome assembly (since long-read genome assemblies are more likely to assemble across repetitive regions), filtering for alignments with >85% identity.

Using this approach, we then determined centromere locations in the reference genome (converting alignment positions in the PacBio genomes to their corresponding or closest reference genome coordinates). To further explore the predicted centromeres, we created dotplots for contigs from the PacBio genomes that spanned a predicted centromere. To do so, we used *nucmer* with --maxmatch, -l 50, -c 100 to align each contig to itself and then plotted all alignments >100 bp using R.

### Characterizing repeat content at inversion breakpoints

#### Dotplots

To evaluate whether inversion breakpoints occurred in repetitive regions, we created dotplots from the PacBio contig-level assemblies. We performed self-v-self nucmer alignments for contigs spanning inversion breakpoints, with --maxmatch, -l 50, -c 100; we filtered for alignments >1 kb and plotted the alignments in R.

#### Inverted repeats and segmental duplications

We identified inverted repeats and segmental duplications (SDs) near inversion breakpoints using the package SEDEF^28^. For the relevant PacBio contigs identified above (spanning or adjacent to inversion breakpoints), we softmasked common repeats with RepeatMasker, using --xsmall and --species rodentia, and masked the 344-bp centromere satellite sequence. We then performed SEDEF with default parameters on the entire set of relevant PacBio contigs. First, we determined inverted repeats to be any repeat identified by SEDEF that mapped in opposite orientation to within 500 kb of both inversion breakpoints. Next, we called repeats as SDs if they were duplicated within 500 kb of a breakpoint, were ≥1 kb in length, had ≥70% identity with a duplication, and had < 70% of its sequence masked as common repeats. We then determined the density of SDs within 500 kb of each inversion breakpoint (note that we excluded breakpoints at chromosome ends since telomeres are not fully assembled in these genome assemblies). To compare the breakpoint SD-density to random regions genome-wide, we also ran SEDEF on each contig from the *P. m. bairdii* PacBio genome assembly and called SDs. We then randomly selected 10,000 sites from across the genome and calculated the density of SDs within 500 kb of each site. Finally, we tested whether inversion breakpoints were significantly enriched for SDs relative to the 10,000 randomized regions using the Kolmogorov-Smirnov test in R.

#### Genes near inversion breakpoints

We used the *P. m. bairdii* genome annotation (Pman2.1_chr_NCBI.corrected.merged-with-Apollo.Aug19.sorted_chr15.gff3) to explore whether inversion breakpoints disrupted annotated protein-coding genes. We tested whether the number of breakpoints disrupting gene sequence was expected by chance based on overall gene density using a binomial test; we calculated the gene density (including exons, introns and UTRs) to be 39% genome-wide and then used *binom*.*test* in R to perform a binomial test, with probability of success=0.39.

### Inversion frequencies

#### Sampling populations across species range

To characterize the frequencies and distributions of the inversions across the *P. maniculatus* range, we included 3 - 46 individuals from each of an additional eight populations, which when combined with the initial populations, yielded a total of 218 mice from 13 populations. For five of the new eight populations (populations *a, b, f, i* and *k* from Figure 4A; see Table S3 for sample details), we extracted DNA from liver tissue and performed whole-genome re-sequencing (∼10-15X coverage) and variant calling as described above. We additionally performed whole-genome re-sequencing for 11 *P. leucopus* samples and 2 *P. californicus* samples (see Table S3 for sample details), which we also included in our variant calling pipeline. For three additional populations (populations *d, g*, and *h* from Figure 4A; see Table S3 for sample details), we obtained publicly available exome-sequencing data^58^ (NCBI: PRJNA528923) and mapped sequencing reads to the *P. maniculatus* reference genome with BWA-MEM. We then performed variant calling as described above, except that these samples were joint-genotyped separately from the whole-genome re-sequenced samples.

#### Phylogenetic trees

To reconstruct the evolutionary relationships among populations, we used RAxML^59^ to build maximum-likelihood trees. First, we created a tree for the five focal *P. maniculatus* populations and two outgroups (*P. leucopus* and *P. californicus*). Using hard-filtered SNPs from across the autosomes, we thinned SNPs to at most 1 SNP per 100 kb using vcftools and merged vcfs across chromosomes. We converted the merged vcf to a PHYLIP matrix using vcf2phylip.py (https://github.com/edgardomortiz/vcf2phylip) and removed invariant sites using ascbias.py (https://github.com/btmartin721/raxml_ascbias), resulting in a total of 12,292 SNPs. We then ran RAxML v8.2.12 using the ASC_GTRCAT model with the conditional likelihood method, -asc-corr=lewis, to correct for the ascertainment bias due to using SNPs^60^. We ran 100 bootstraps, with “-f a” to perform rapid bootstrap analysis and visualized trees in iTOL^61^. We next created a tree for all 13 *P. maniculatus* populations and the two outgroups. To do so, we first merged the variants called for the three exome-sequenced populations with the whole-genome re-sequenced vcfs and subset each population to at most 15 individuals. We removed variants with missing genotypes for >20% of samples and masked inversion regions using bcftools. We then converted the vcf to a PHYLIP matrix and removed invariant sites as described above, resulting in a total of 15,518 SNPs. We ran 100 bootstraps, with “-f a” to perform rapid bootstrap analysis and visualized trees in iTOL.

#### Genotyping samples for inversions

To genotype individuals for the presence/absence of inversions, we used a PCA approach. For each inversion, we selected closely related populations segregating for the inversion of interest and performed PCA for that inversion region using *scikit-allel*, as described above. Performing PCA with only a subset of populations allowed for the inversion of interest (rather than population divergence) to drive variance along PC1. We then projected the remaining samples onto the PC1 and PC2 axes. We genotyped samples for each inversion based on loading scores along PC1 (along which samples clustered into inversion genotype groups) with manual determination of boundaries. We verified that samples called as inversion heterozygotes had elevated heterozygosity in the inversion region using the *count_het* function in *scikit*-*allel*. We set any populations with ambiguous clustering along PC1 for a given inversion to have missing genotypes. Finally, we determined inversion genotype frequencies for each population and tested for deviations from HWE using *HWE*.*chisq* in R from the *genetics* package.

We also determined inversion genotypes for (1) 547 F2 hybrids from the *P. m. rubidus* x *P. m. gambelii* cross, and (2) the 136 wild-caught mice from the environmental transect. To do so, we first created a set of SNPs fixed between the inversion and standard arrangements using homozygous samples from only forest (*P. m. rubidus*) and prairie (*P. m. gambelii*) populations, unless there were fewer than three homozygous samples per genotype, in which case we included additional homozygous samples from nearby populations (populations *b* and *f*, Figure 4A) to improve filtering. Previously, the F2 hybrids were sequenced using the ddRAD-sequencing pipeline (as described in section ‘Recombination rates’, NCBI: PRJNA687993) and the 136 transect mice were whole-genome re-sequenced at low coverage (NCBI: PRJNA688305)^19^. Using these sequencing data, we selected the fixed inversion-standard SNPs from bam files for the F2 hybrids and transect mice using *mpileup* and performed the hidden Markov model step of the multiplexed shotgun genotyping pipeline^62^ to determine genotype for each inversion.

#### Mutational load

To test whether the inversions were enriched for deleterious mutations compared to standard haplotypes, we analyzed the number of segregating non-synonymous (*pN*) versus synonymous (*pS*) sites and nucleotide diversity at non-synonymous (*π*_*N*_) versus synonymous (*π*_*S*_) sites using *PopGenome*^63^. For each inversion, we selected samples homozygous for the inversion arrangement and used *readVCF* to import biallelic SNPs for the samples and inversion region of interest into *PopGenome*; specifically, we selected homozygous samples from the major *P. maniculatus* clade (Figure 4B; populations *a, b, c, e, f, i* and *j*) except for inv10.0 and inv11.0, for which we also included populations *k, l, m* in order to sample both homozygous genotypes. We then used the *set*.*synnonsyn* function with the *P. m. bairdii* genome annotation to determine non-synonymous and synonymous sites. Next, we computed nucleotide diversity for each synonymous and non-synonymous site with the *diversity*.*stats* function. Finally, for 500-kb windows across each inversion region, we calculated the ratio of segregating non-synonymous versus synonymous sites (*pN*/*pS*) and the ratio of mean nucleotide diversities for non-synonymous versus synonymous sites (*π*_*N*_/*π*_*S*_) (using only sites that were segregating within the homozygous sample set). We then repeated these analyses for samples homozygous for the standard arrangement. To test whether the inversion and standard haplotypes significantly differed in *pN*/*pS* or *π*_*N*_/*π*_*S*_, we performed two-sided t-tests in R. Inv7.1 was excluded from this analysis because we had sequencing data for only one homozygous inversion sample; inv9.1 was excluded from this analysis because it harbors only six genes.

#### SLiM simulations

To explore a possible role of selection on the inversions, we performed forward-genetic simulations in SLiM v3.6^33^. We simulated the forest (population *c, P. m. rubidus*) and prairie (population *e, P. m. gambelii*) populations evolving under a previously estimated best-fit demographic model^19^ and introduced an inversion as a Mendelian locus as a single copy. We set separate selection coefficients for the inversion locus in the forest versus prairie populations, varying the selection coefficients from -0.01 to +0.01. We introduced the inversion into either the forest or prairie population at five timepoints, corresponding to 1.5e4, 1.5e5, 7.5e5, 1.5e6, and 2.2e6 generations ago, with 2.2e6 being the estimated time of the forest-prairie split. To reduce computational time, we scaled parameters by a factor of 100, with population sizes and times divided by 100 (e.g., after scaling, timepoints ranged from 1.5e2 – 2.2e4 generations ago) and migration rates and selection coefficients multiplied by 100 (e.g., after scaling, selection coefficients ranged from -1.0 to +1.0), to keep *Nm* and *Ns* consistent^33^. For each set of forest and prairie selection coefficients and each timepoint, we ran 1,000 simulations and recorded the frequency of the inversion in the forest and prairie populations at the end of the simulation. Finally, for each scenario, we computed the probability that the inversion reached an absolute allele frequency difference between the forest and prairie populations >50%. All selection coefficients are reported as their values before scaling.

#### Clinal variation

To test whether inversion frequency is associated with local habitat, we analyzed *P. maniculatus* mice previously collected across a forest-prairie environmental gradient, which includes 136 samples from 9 sites across the Cascade mountains in Oregon, USA^19^. Using publicly available sequencing data^19^ (NCBI: PRJNA688305), we genotyped the 136 samples for the inversions (see ‘Genotyping samples for inversions’ section above) and then used the package HZAR v0.2.5^64^ to fit clines to inversion genotypes (https://github.com/oharring/chr15_inversion). We fit ten different cline models, by varying scaling of minimum and maximum allele frequencies (scaling = ‘fixed’ or ‘free’) and how exponential tails were fit (tails = ‘none’, ‘left’, ‘right’, ‘mirror’, and ‘both’). We selected the best model for each inversion using AICc values. Clines shown in Figure 6B are fit with tails=‘none’ and scales=‘fixed’; best-fit clines are shown in Figure S7.

#### Genotype-phenotype associations

Using data from a reciprocal intercross between *P. m. rubidus* (forest population) x *P. m. gambelii* (prairie population) F2 hybrids (*n* = 547) described above, we tested for associations between inversion genotype and three forest-ecotype defining traits: tail length, foot length and coat color. We used previously published phenotypic measurements^19^ and the inversion genotypes reported here. For each of the 13 polymorphic forest-prairie inversions, we tested whether inversion genotype was significantly correlated with trait variation using linear models in R, with genotype coded numerically (additive genetic model); for tail and foot length, we included body length as a fixed effect. We corrected for multiple hypothesis testing (i.e., testing 13 different inversions) using Bonferroni correction.

## Acknowledgments

We thank T. Sackton, D. Khost and members of the Hoekstra lab for their advice on the analyses; T. Sackton, J. Mallet, L. Gozashti, A. Kautt, and members of the Mallet lab for providing helpful feedback on the manuscript; T.B. Wooldridge for sharing short-read sequencing data; and E. Hager and T.B. Wooldridge for many helpful discussions on inversions. The Bauer Core Facility at Harvard University provided short-read library preparation and sequencing services. The University of Washington PacBio Sequencing Core provided long-read library preparation and sequencing services. Computational analyses were run on the Odyssey and Cannon clusters supported by the Faculty of Arts and Sciences Research Computing Group at Harvard University. We thank the Museum of Southwestern Biology (University of New Mexico), Museum of Comparative Zoology (Harvard University), Sam Cushman (US Forest Service, Rocky Mountain Research Station), and Cody Thompson (University of Michigan) for providing specimens used in this study.

## Funding

OSH was supported by a National Science Foundation (NSF) Graduate Research Fellowship, a Harvard Quantitative Biology Student Fellowship (DMS 1764269), the Molecular Biophysics Training Grant (NIH NIGMS T32GM008313), an American Society of Mammalogists Grants-in-Aid of Research, and a Society for the Study of Evolution R.C. Lewontin Early Award. HEH is an Investigator of the Howard Hughes Medical Institute.

## Author contributions

OSH conceived of the study and performed the analyses, with input from HEH. OSH and HEH wrote the manuscript.

## Competing interests

Authors declare no competing interests.

## Data and materials availability

Associated data and code will be uploaded to NCBI SRA and GitHub before publication.

## Permissions and approval

All experiments were approved by Harvard University’s IACUC.

## Supplementary Materials for

**Figure S1.**
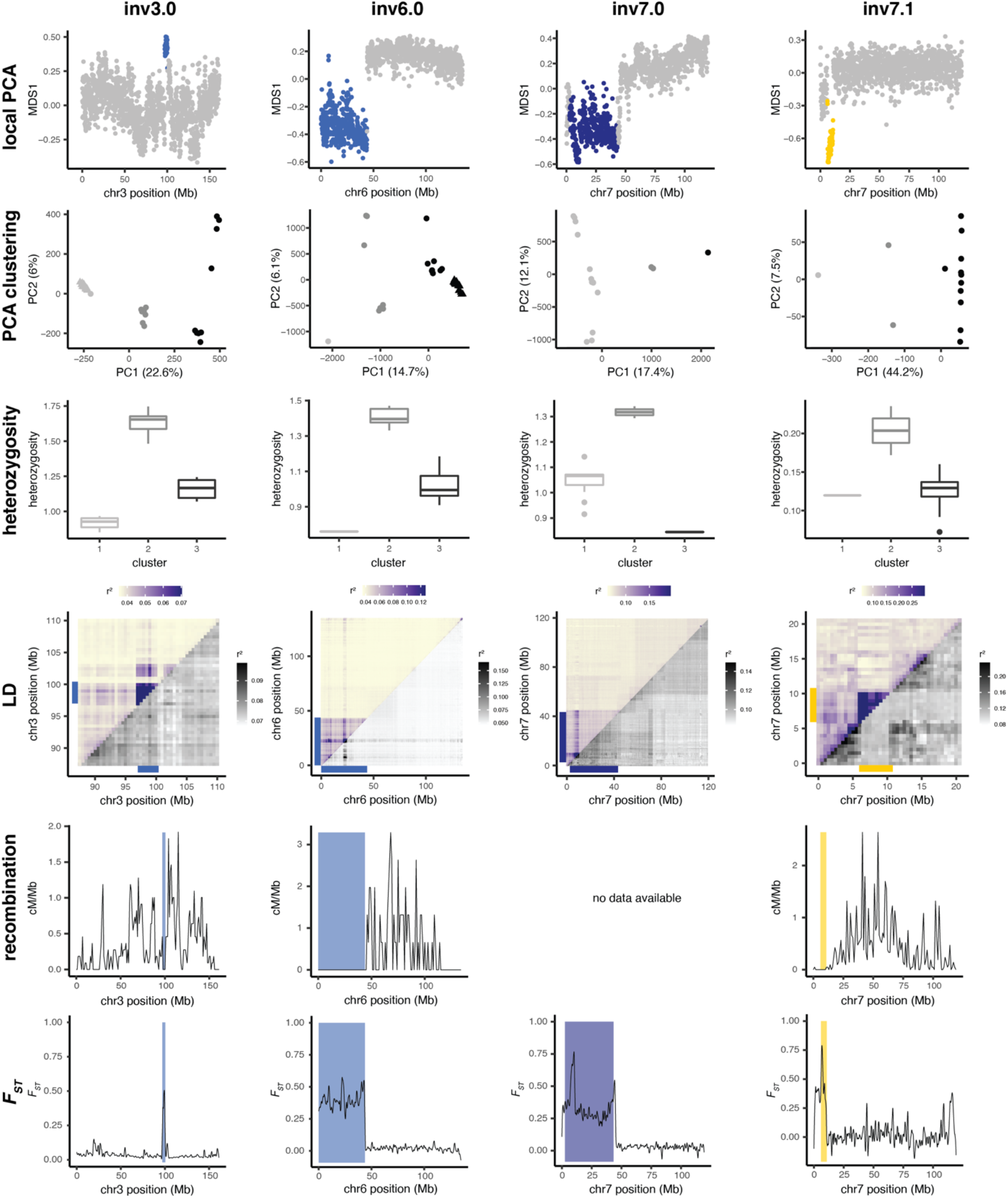
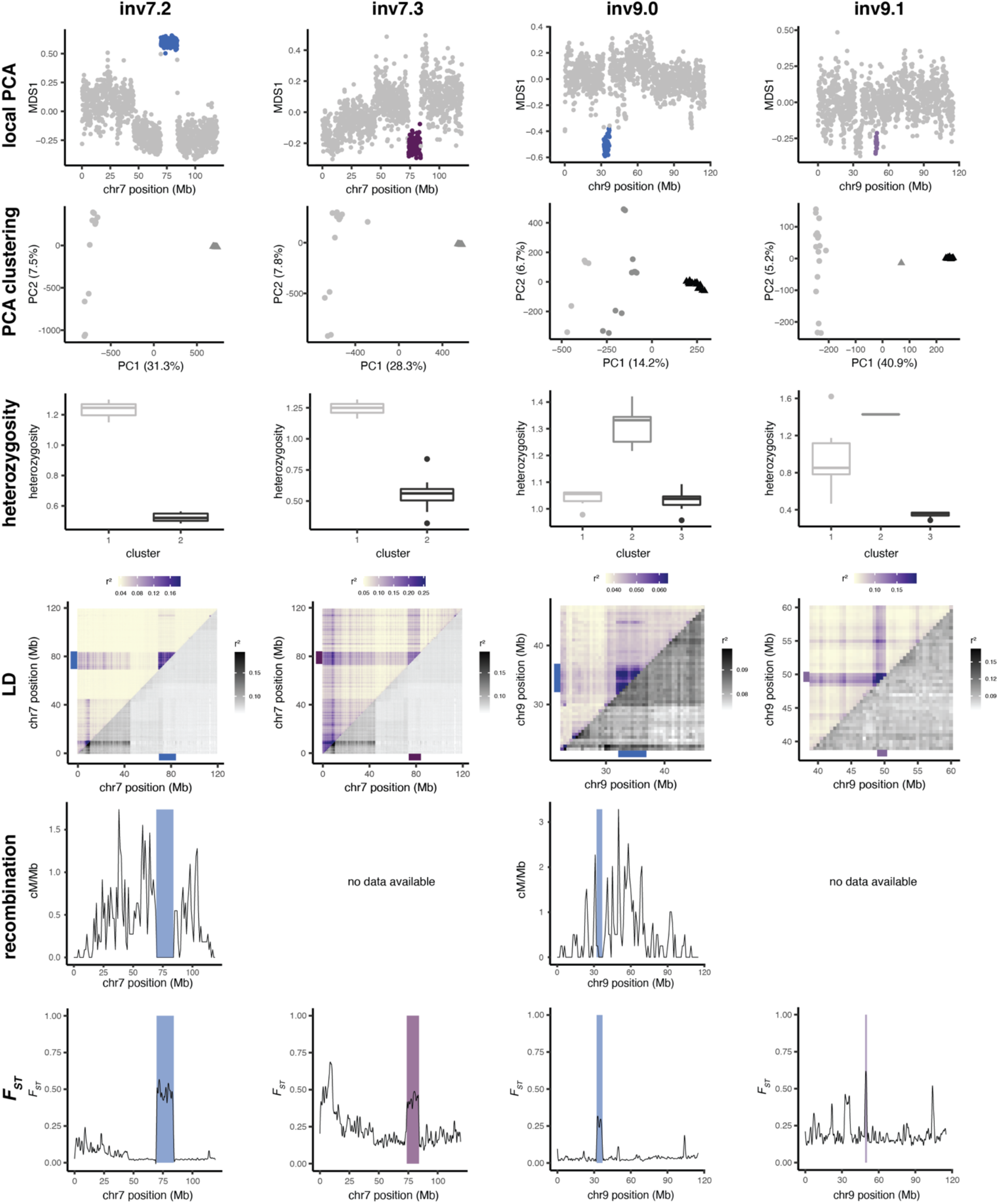
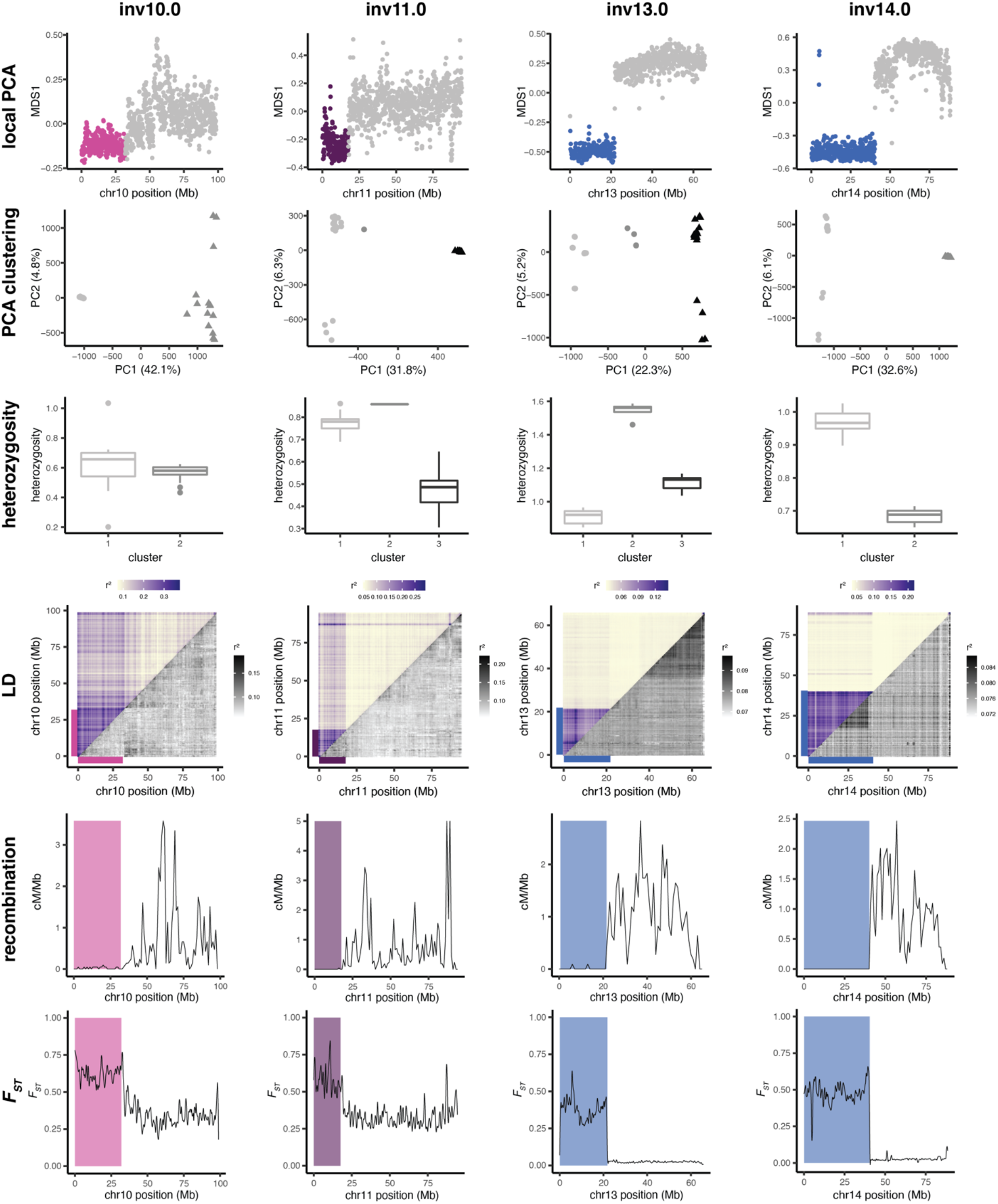
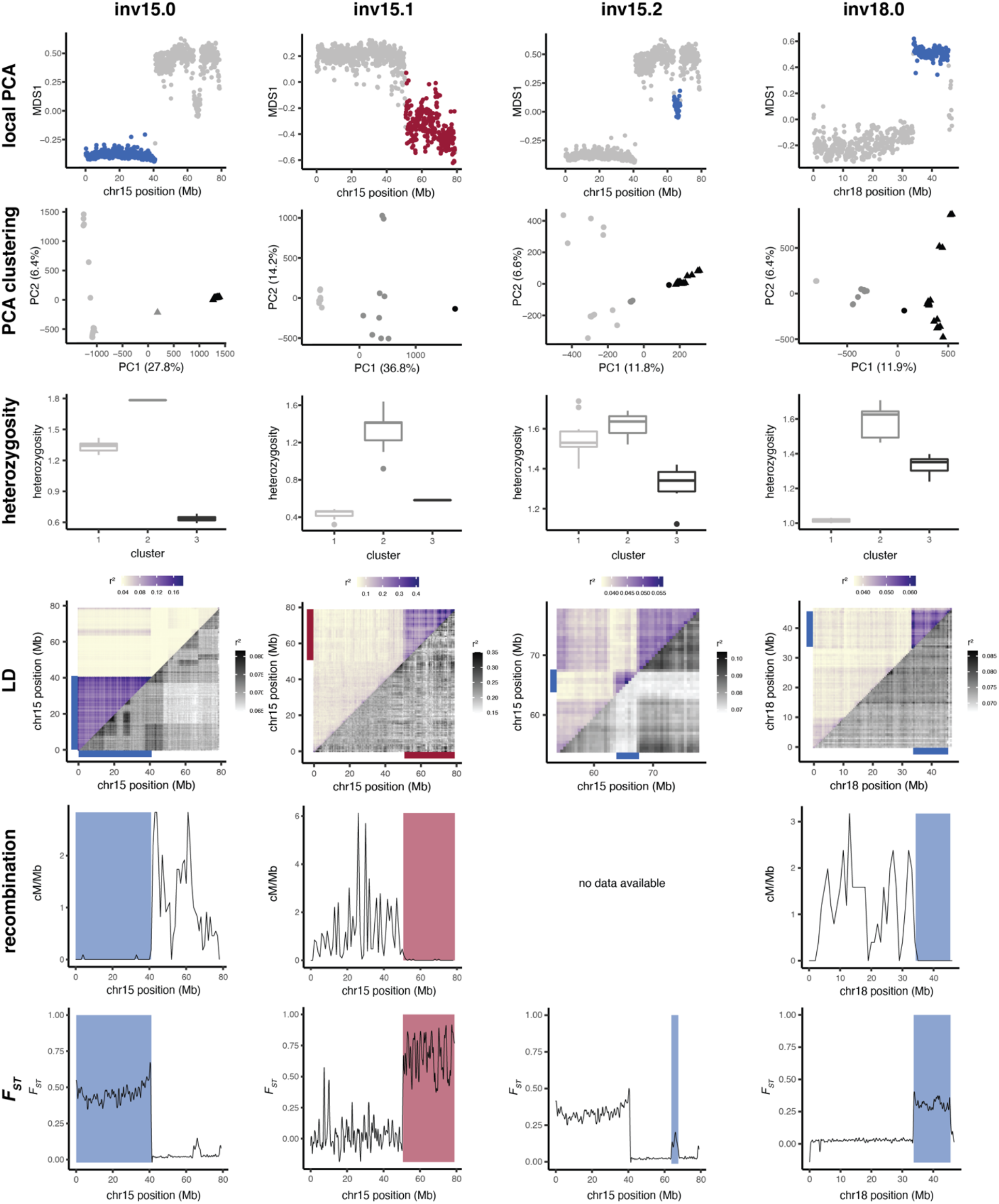
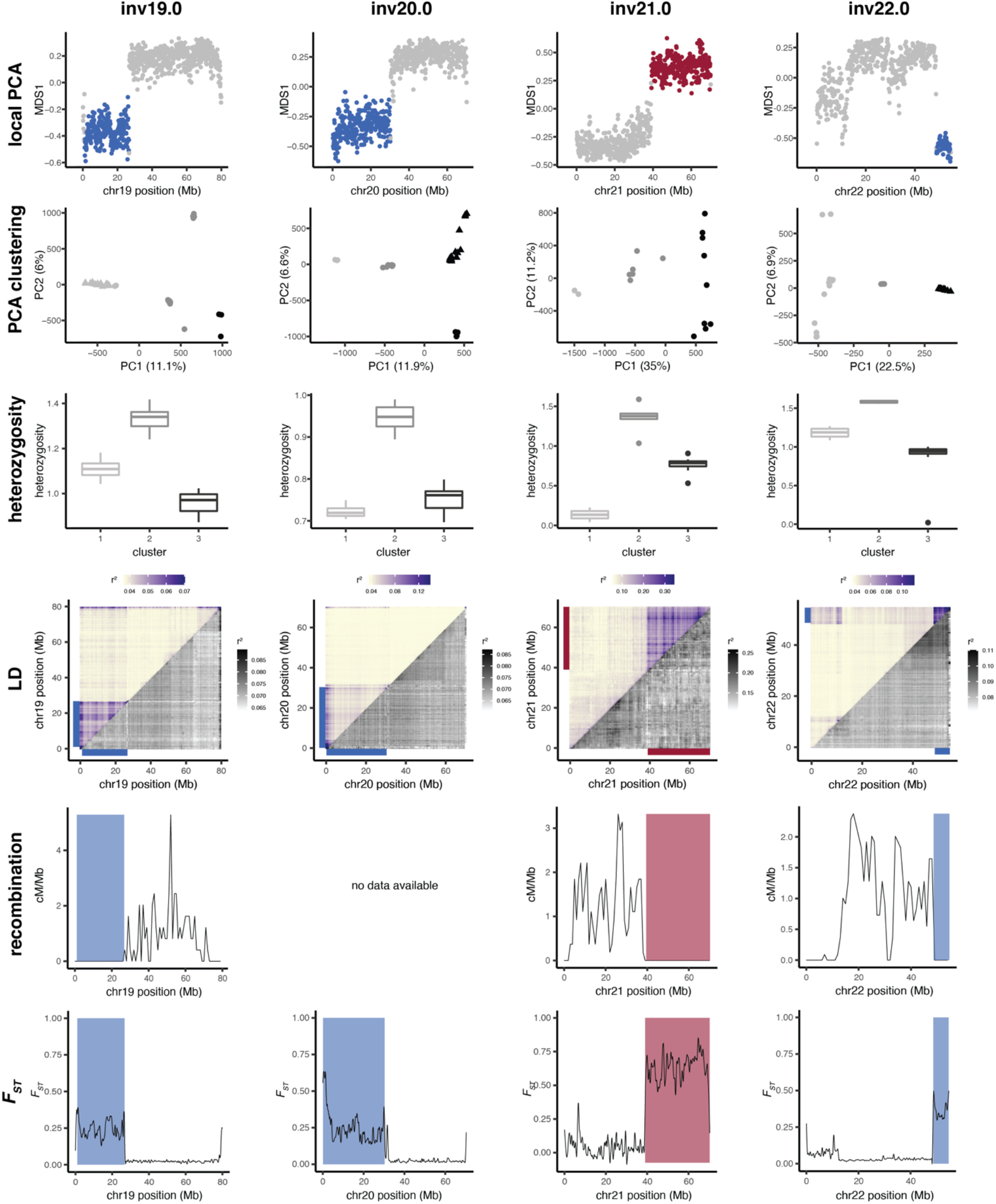
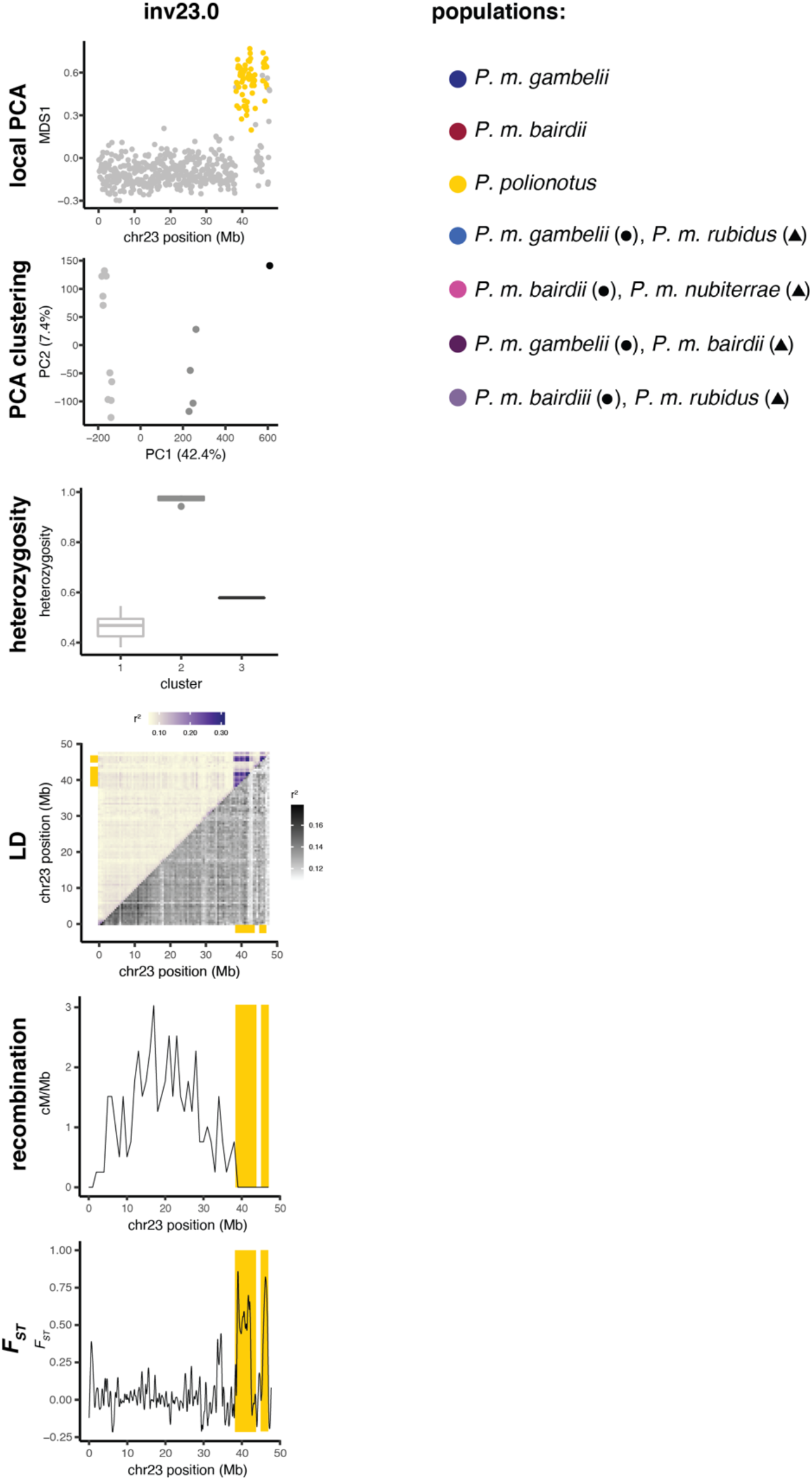
Identifying inversion polymorphisms based on population genomic signatures. For each identified inversion polymorphism, the following signatures of inversions are shown (colors correspond to focal population or population-pair in which inversion was identified, see legend): (1) Local PCA performed with *lostruct*, where each dot represents a 100-kb window. Distances between local PCA maps are represented by the MDS1 axis, with outlier windows highlighted in color. (2) Clustering of samples by PCA for entire outlier region found with local PCA, assigned using k-means clustering. (3) Heterozygosity (percent of sites that are heterozygous) of outlier region for samples by cluster assignments from PCA above. Boxplots indicate upper and lower quartiles, with median (center line); whiskers show 1.5x interquartile range; points show outliers. (4) LD for chromosomes harboring the example inversions, shown as mean *r*^2^ values for paired windows across each chromosome. Upper triangle shows mean *r*^2^ values including all samples from PCA clustering. Lower triangle shows mean *r*^2^ values for only the more common homozygote genotype as determined in PCA clustering. Colored bars highlight outlier region from local PCA. Scales for *r*^2^ values provided. (5) Recombination rates in cM/Mb shown for lab-born inversion heterozygotes. Outlier region found in local PCA is highlighted. Five inversions have missing data since inversion heterozygotes were not measured in the lab. (6) *F*_*ST*_ between homozygous genotypes (clusters 1 and 3 from PCA and heterozygosity plots). Outlier regions found in local PCA are highlighted. Note that the discontinuity for inv23.0 is likely due to reference genome mis-assembly.

**Figure S2.**
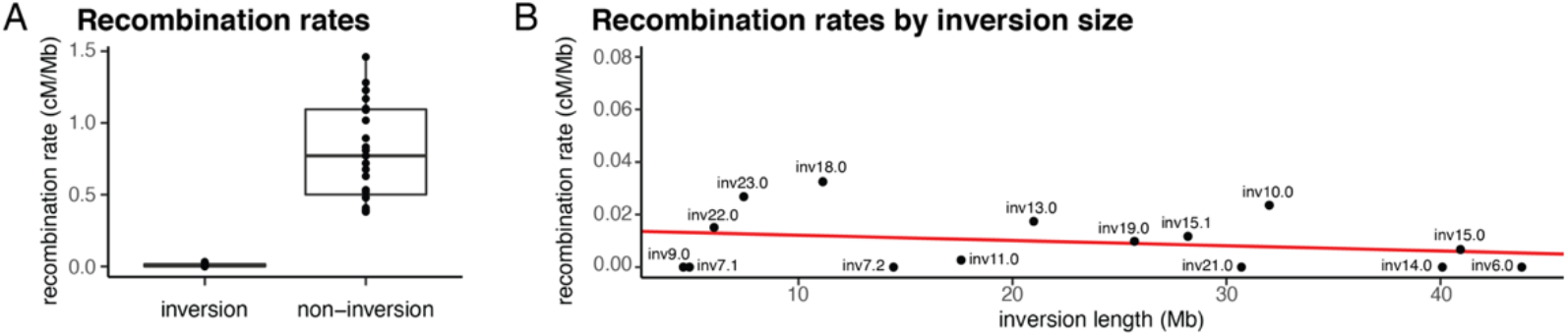
Recombination effects of inversion heterozygotes. (**A**) Recombination rates for inversion versus non-inversion regions from lab-born F_2_ hybrids. Recombination rates for inversion regions are measured in inversion heterozygotes only; rates for non-inversion regions include all lab-born F_2_ hybrids. Boxplots indicate upper and lower quartiles, with median (center line); whiskers show 1.5x interquartile range. Points for inversion regions represent inversions; points for non-inversion regions represent chromosomes (excluding inversion regions). Recombination rate for inversion regions = 0.01±0.03; non-inversion regions = 0.80±0.34 (mean±sd). (**B**) Recombination rates for inversion regions by inversion size in megabases. Linear fit (red line, p>0.05).

**Figure S3.**
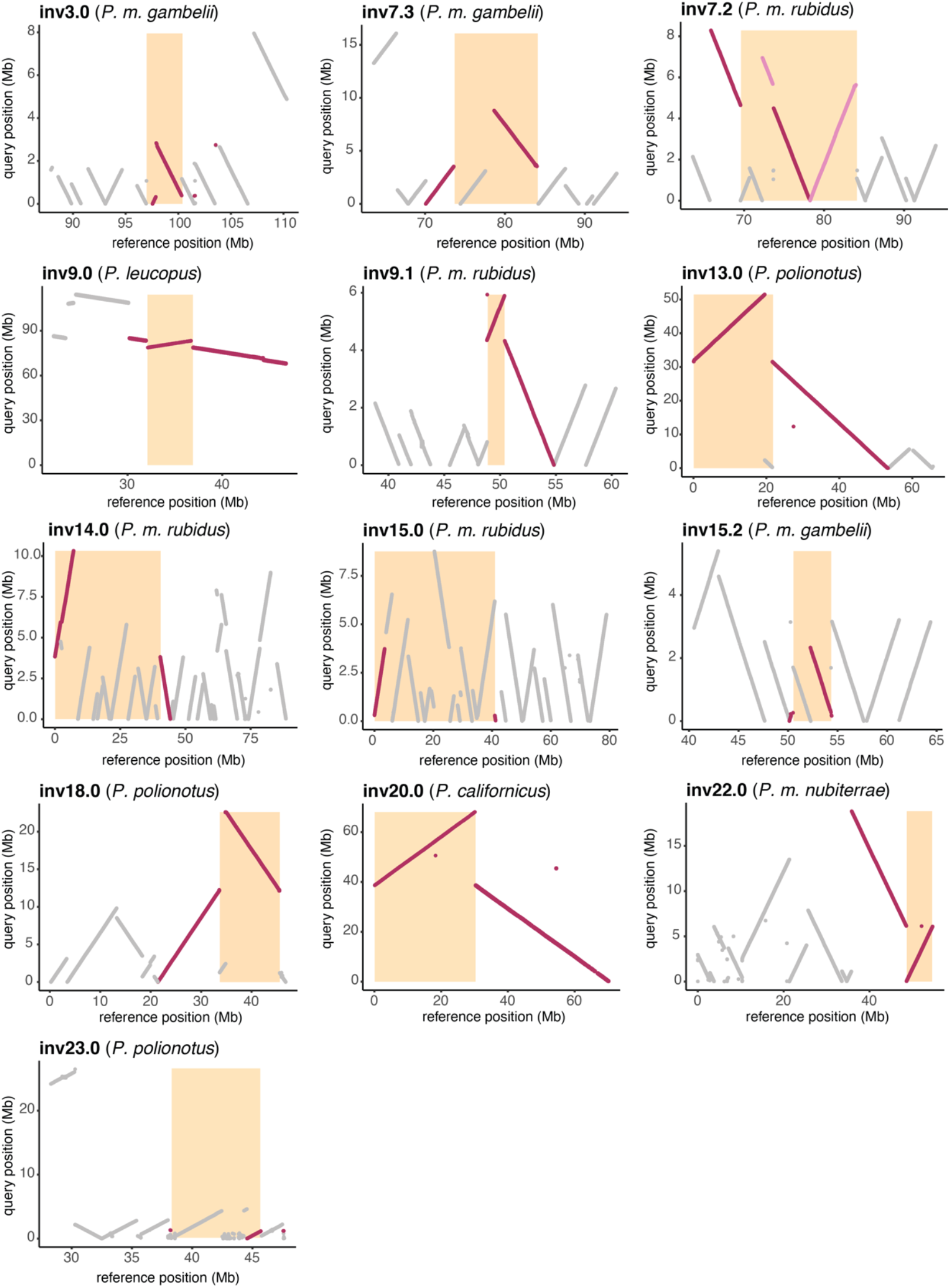
Localizing inversion breakpoints. Contigs highlighting breakpoints for 13 inversions. Contigs from *de novo* genome assemblies (‘query’, y-axis) were aligned to the *P. maniculatus* reference genome (‘reference’, x-axis) with *nucmer*. Populations corresponding to the *de novo* assembly used in each plot are given; for inv9.0 and inv20.0 inversions, breakpoints were localized by aligning *P. leucopus* and *P. californicus* reference genomes to the *P. maniculatus* reference genome, respectively. Contigs (gray) and those identifying inversion breakpoints (red) are shown. Predicted inversion boundaries are highlighted (orange box). For the inv7.2 plot, the pink contig highlights a derived inversion (inv7.3) in the reference genome; when the reference genome is re-oriented to the ancestral state, the contig highlighted in red shows the inv7.2 inversion breakpoints.

**Figure S4.**
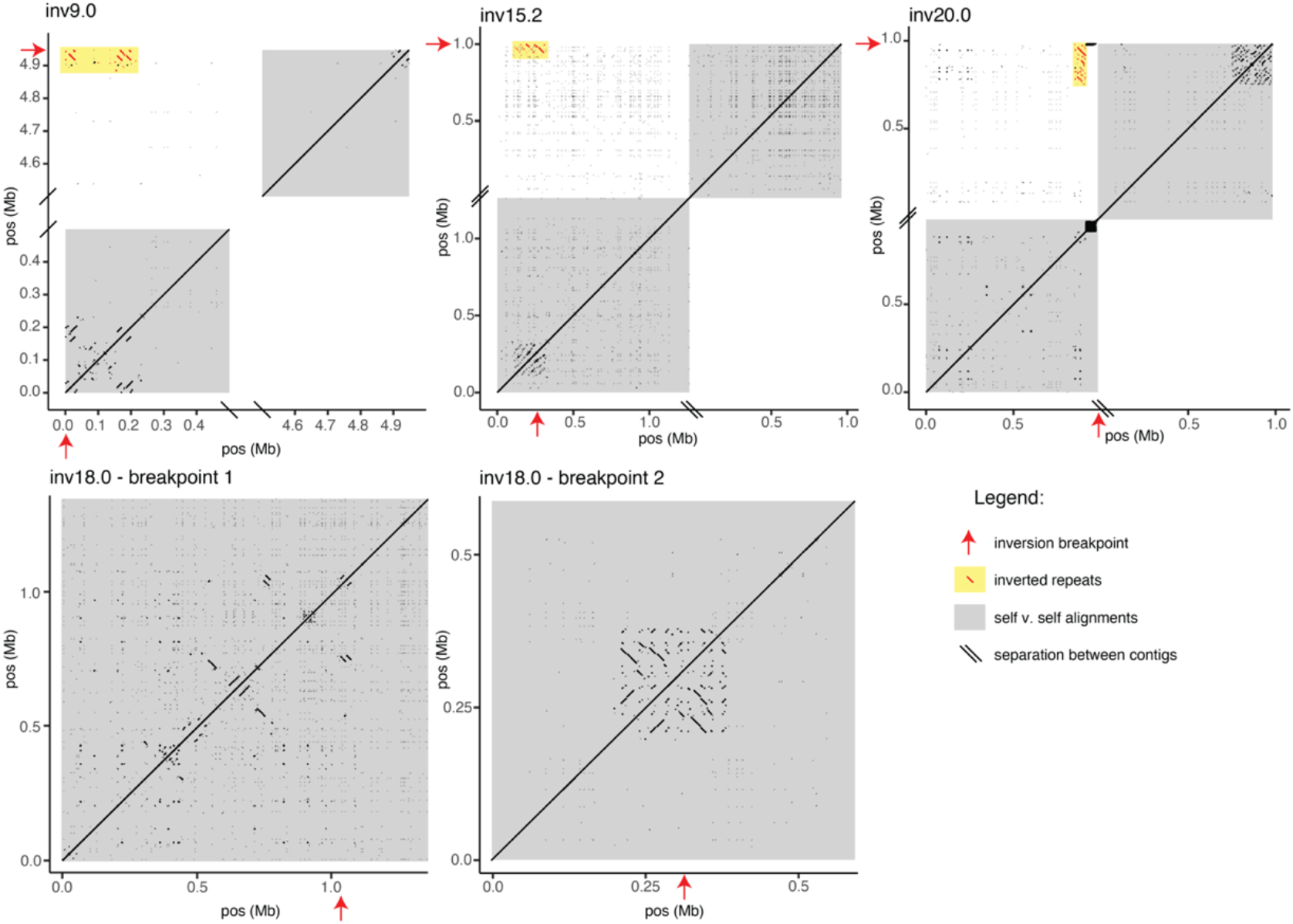
Inverted repeats and segmental duplications. Examples of inversion breakpoints near large inverted repeats (inv9.0, inv15.2, inv20.0) and segmental duplications (inv9.0, inv15.2, inv20.0, inv18.0). Dotplots show alignments for long-read assembly contigs spanning or nearly spanning breakpoints. Self-v-self alignments are highlighted (gray boxes), with alignments between breakpoint regions (within 500 kb of breakpoints) shown in upper left quadrant for inv9.0, inv15.2 and inv20.0. Location of breakpoints (red arrows) shown; only alignments with length >100 bp and within 500 kb of the breakpoints are shown. Inverted repeats mapping to within 500 kb of both breakpoints are shown (red) and highlighted (yellow boxes). Self-v-self alignments also show segmental duplications near breakpoints.

**Figure S5.**
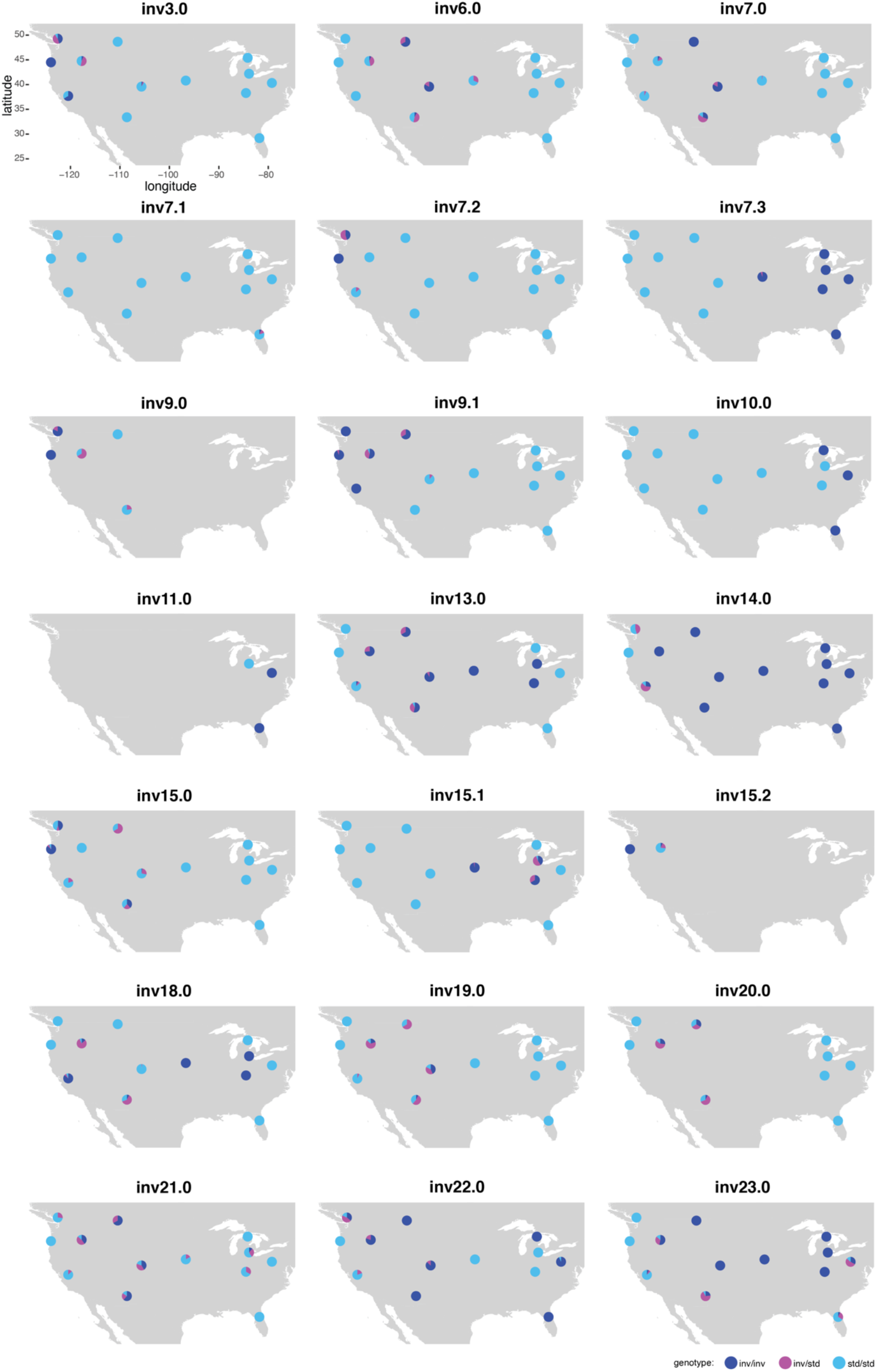
Distributions of inversions across species range. Genotype frequencies shown across species range for each inversion (for 13 populations shown in Figure 4A). Inversions were genotyped with PCA; populations with ambiguous genotypes for a given inversion are not included.

**Figure S6.**
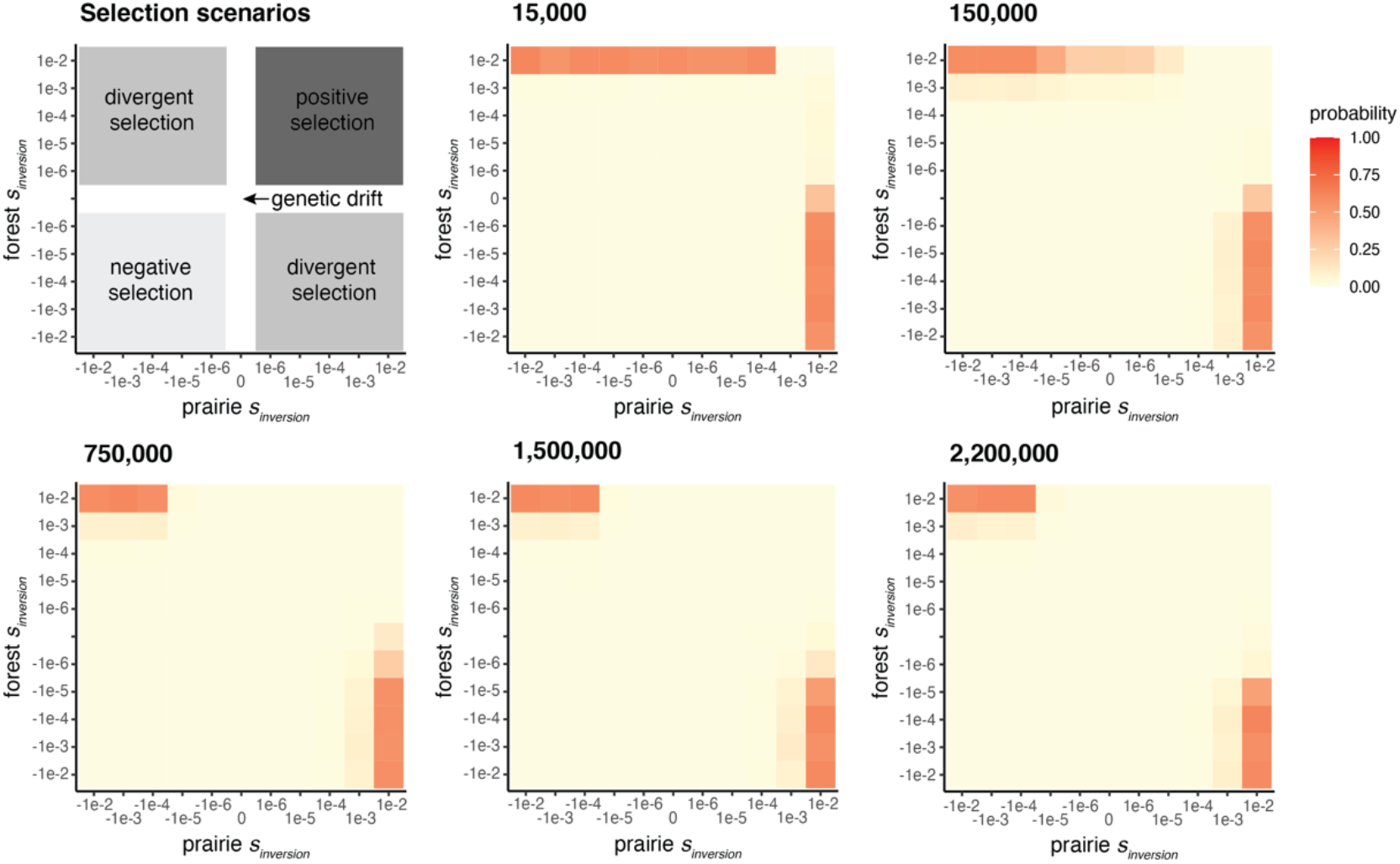
Forward genetic simulations of selection on inversions. The evolution of an inversion was simulated in SLiM under a best-fitting demographic model of the forest and prairie populations. The inversion locus was introduced into the populations at five timepoints (shown in generations ago); the final timepoint (2.2m generations) represents the forest-prairie split time estimate. Simulations for a range of selection coefficients for the inversion were performed, with 1,000 simulations per scenario. The selection scenarios are shown (upper left). Heatmaps show the probability of the inversion reaching a forest-prairie allele frequency difference >50% for each combination of selection coefficients.

**Figure S7.**
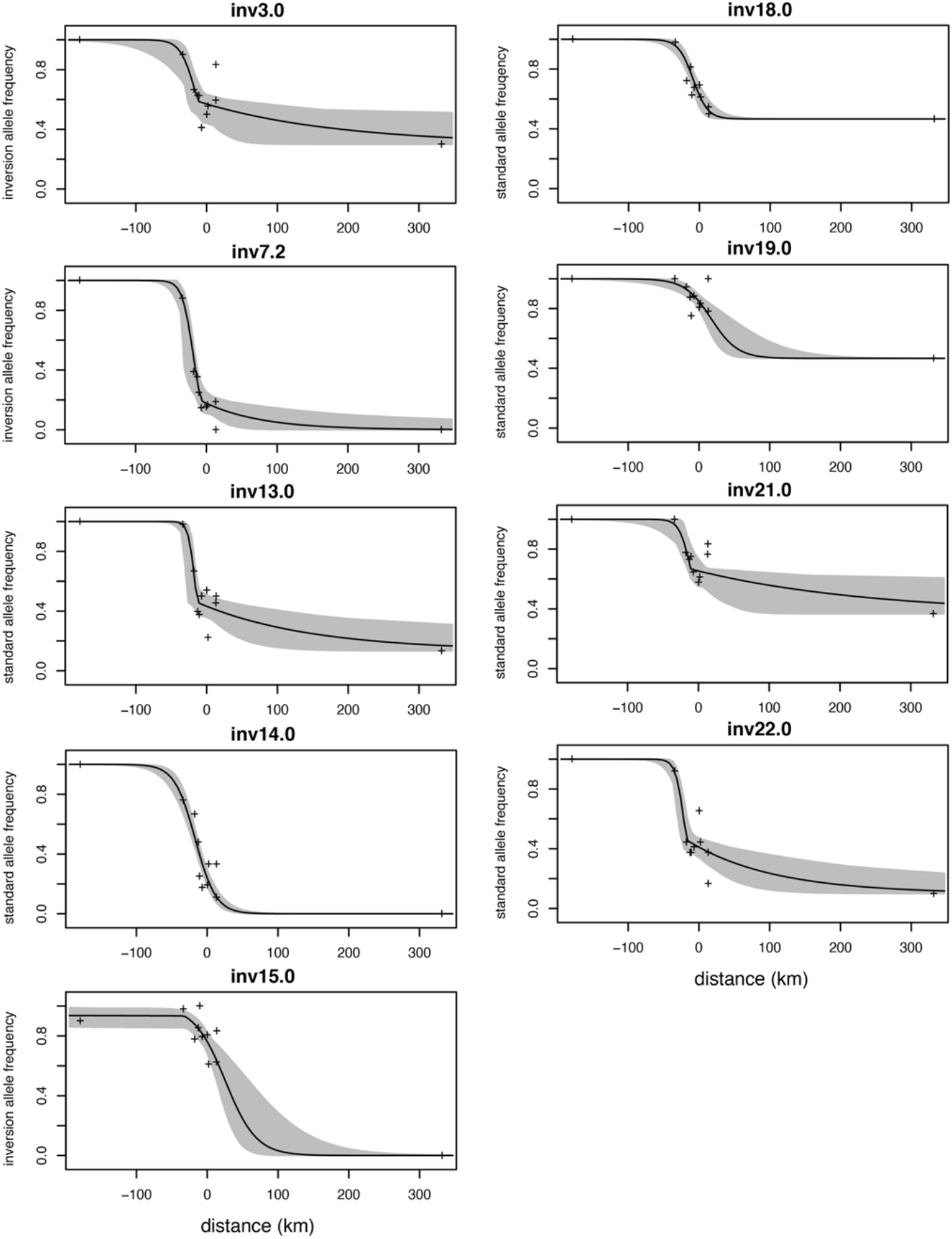
Clinal variation in inversion frequencies. Inversion genotype frequencies shown across an environmental transect, with clines fit using *hzar*. Best-fit clines with 95% credible cline region shown. Sampled populations are highlighted (black crosses), with focal forest (left-most) and prairie (right-most) populations. Major allele in forest population is plotted, with y-axis label indicating *inversion* or *standard* haplotype.

**Table S1.** Summary statistics for *de novo* genome assemblies with PacBio long-read sequencing. Contig-level genomes were assembled using *flye* for one individual from each of the five focal populations. PacBio sequencing outputs are reported as subread N50 and *flye* genome assemblies are summarized with total length of assemblies, contig N50s, and number of contigs. Differences in assembly contiguity are likely driven by differences in heterozygosity in the sequenced samples.

| **Population** | **PacBio**<br>**subread**<br>**N50 (kb)** | **Flye**<br>**assembly**<br>**total length**<br>**(Gb)** | **Flye**<br>**assembly**<br>**contig**<br>**N50 (Mb)** | **Flye**<br>**assembly**<br>**# of**<br>**contigs** |
| --- | --- | --- | --- | --- |
| *P. maniculatus bairdii* | 36.9 | 2.53 | 11.2 | 5,747 |
| *P. maniculatus gambelii* | 36.6 | 2.60 | 3.2 | 12,590 |
| *P. maniculatus rubidus* | 37.9 | 2.61 | 2.6 | 15,237 |
| *P. maniculatus nubiterrae* | 40.2 | 2.56 | 5.5 | 8,512 |
| *P. p. subgriseus* | 38.8 | 2.51 | 9.5 | 3,108 |

**Table S2.**
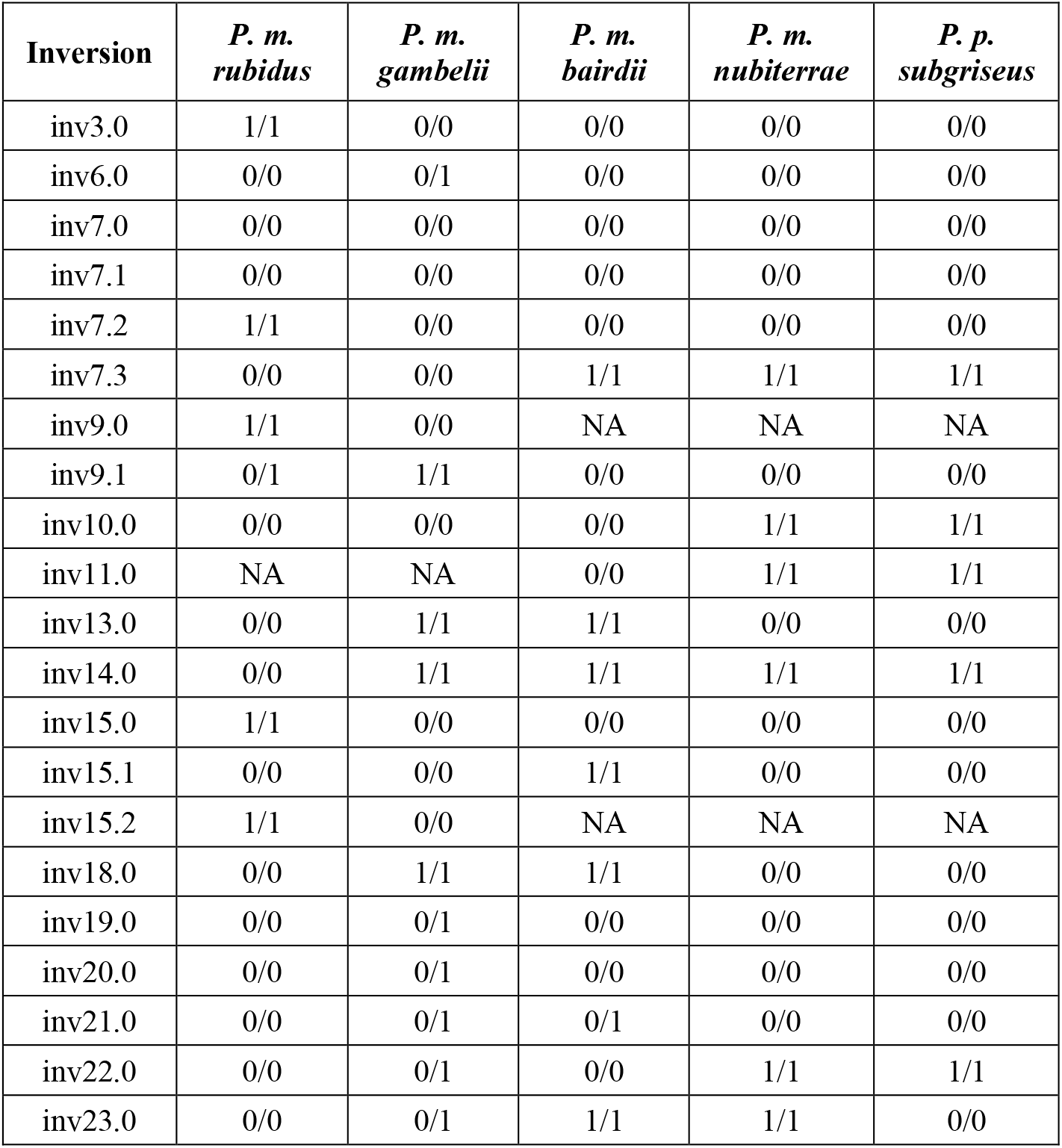
Inversion genotypes for long-read samples. Inversion genotypes (0=standard, 1=inversion) for all 21 inversions in each of the five PacBio long-read sequenced samples. Inv6.0, inv7.0, inv7.1, inv19.0, inv20.0, inv21.0 are not represented by both homozygous genotypes in these samples.

**Table S3.** List of samples for each population included in this study. Samples from populations *a* – *m* included in this study, listed by population from Figure 4A, with subspecies labels for focal populations, and ecotype labels for forest and prairie populations with the following descriptors: *Sample*.*ID*: unique ID for each sample; *Museum*.*ID*: museum accession number for that sample; *Latitude/Longitude*: geographic coordinates from which sample was collected; *Source*: institution, researcher or NCBI project from which the tissue or sequencing data was obtained (MCZ = Museum of Comparative Zoology, Harvard University; MSB = Museum of Southwestern Biology, University of New Mexico; USFS = US Forest Service; UM = University of Michigan; internal = internal from Hoekstra lab); *Sequencing*: type of sequencing data obtained for that sample (WGS = whole-genome re-sequencing; exome = exome-sequencing); *SRA*.*run*: NCBI SRA run identifier.

| Sample.ID | Museum.ID | Latitude | Longitude | Source | Sequencing | SRA.run |
| --- | --- | --- | --- | --- | --- | --- |
| **Population a** |  |  |  |  |  |  |
| FSB11 | MCZ69737 | 49.28 | -122.57 | MCZ | WGS |  |
| FSB25 | MCZ69784 | 49.27 | -122.57 | MCZ | WGS |  |
| FSB37 |  | 49.28 | -122.57 | internal | WGS |  |
| FSB42 |  | 49.28 | -122.57 | internal | WGS |  |
| FSB43 | MCZ69794 | 49.28 | -122.57 | MCZ | WGS |  |
| FSB48 | MCZ69756 | 49.27 | -122.57 | MCZ | WGS |  |
| FSB6 | MCZ69733 | 49.27 | -122.57 | MCZ | WGS |  |
| FSB7 | MCZ69734 | 49.27 | -122.57 | MCZ | WGS |  |
| FSB51 |  | 49.27 | -122.58 | internal | WGS |  |
| FSB52 | MCZ69797 | 49.27 | -122.58 | MCZ | WGS |  |
| FSB53 |  | 49.27 | -122.58 | internal | WGS |  |
| **Population b** |  |  |  |  |  |  |
| 476_3_1 |  | 48.66 | -110.46 | Sam Cushman (USFS) | WGS |  |
| 477_1_4 |  | 48.66 | -110.46 | Sam Cushman (USFS) | WGS |  |
| 479_2_1 |  | 48.66 | -110.46 | Sam Cushman (USFS) | WGS |  |
| **Population c (*P. m. rubidus*; forest ecotype)** |  |  |  |  |  |  |
| ERH21 | MCZ69553 | 44.24 | -123.95 | PRJNA688305 | WGS | SRR13318612 |
| ERH27 | MCZ69544 | 44.27 | -124.04 | PRJNA688305 | WGS | SRR13318609 |
| ERH36 | MCZ69566 | 44.22 | -124.01 | PRJNA688305 | WGS | SRR13318608 |
| ERH52 | MCZ69555 | 44.24 | -123.95 | PRJNA688305 | WGS | SRR13318601 |
| ERH19 | MCZ70244 | 44.27 | -124.03 | PRJNA688305 | WGS | SRR13318613 |
| ERH48 | MCZ69563 | 44.22 | -124.01 | PRJNA688305 | WGS | SRR13318602 |
| ERH26 | MCZ70248 | 44.27 | -124.04 | PRJNA688305 | WGS | SRR13318610 |
| ERH14 | MCZ70242 | 44.27 | -124.03 | PRJNA688305 | WGS | SRR13318614 |
| ERH25 | MCZ70247 | 44.22 | -123.92 | PRJNA688305 | WGS | SRR13318611 |
| ERH38 |  | 44.22 | -124.01 | PRJNA688305 | WGS | SRR13318607 |
| ERH42 | MCZ70254 | 44.27 | -124.03 | PRJNA688305 | WGS | SRR13318606 |
| ERH12 | MCZ70240 | 44.27 | -124.04 | PRJNA688305 | WGS | SRR13318616 |
| ERH43 |  | 44.22 | -124.01 | PRJNA688305 | WGS | SRR13318605 |
| ERH46 | MCZ69562 | 44.22 | -124.01 | PRJNA688305 | WGS | SRR13318603 |
| ERH53 |  | 44.24 | -123.95 | PRJNA688305 | WGS | SRR13318600 |
| **Population d** |  |  |  |  |  |  |
| SRR8782118 | MVZ207860 | 37.55 | -120.36 | PRJNA528923 | exome | SRR8782118 |
| SRR8782119 | MVZ207863 | 37.53 | -120.35 | PRJNA528923 | exome | SRR8782119 |
| SRR8782120 | MVZ207864 | 37.54 | -120.49 | PRJNA528923 | exome | SRR8782120 |
| SRR8782121 | MVZ207843 | 37.62 | -120.52 | PRJNA528923 | exome | SRR8782121 |
| SRR8782123 | MVZ207839 | 37.67 | -120.48 | PRJNA528923 | exome | SRR8782123 |
| SRR8782191 | MVZ207878 | 37.70 | -120.24 | PRJNA528923 | exome | SRR8782191 |
| SRR8782192 | MVZ207847 | 37.51 | -120.40 | PRJNA528923 | exome | SRR8782192 |
| SRR8782195 | MVZ207842 | 37.67 | -120.47 | PRJNA528923 | exome | SRR8782195 |
| SRR8782196 | MVZ207851 | 37.55 | -120.35 | PRJNA528923 | exome | SRR8782196 |
| SRR8782197 | MVZ207866 | 37.63 | -120.21 | PRJNA528923 | exome | SRR8782197 |
| SRR8782200 | MVZ207870 | 37.66 | -120.22 | PRJNA528923 | exome | SRR8782200 |
| SRR8782201 | MVZ207856 | 37.55 | -120.36 | PRJNA528923 | exome | SRR8782201 |
| SRR8782202 | MVZ207828 | 37.50 | -120.07 | PRJNA528923 | exome | SRR8782202 |
| SRR8782205 | MVZ207830 | 37.64 | -120.15 | PRJNA528923 | exome | SRR8782205 |
| SRR8782207 | MVZ207832 | 37.67 | -120.46 | PRJNA528923 | exome | SRR8782207 |
| **Population e (*P. m. gambelii*; prairie ecotype)** |  |  |  |  |  |  |
| ERH108 | MCZ69536 | 44.85 | -117.64 | PRJNA688305 | WGS | SRR13318593 |
| ERH87 | MCZ70287 | 44.81 | -117.66 | PRJNA688305 | WGS | SRR13318653 |
| ERH99 | MCZ70296 | 44.81 | -117.60 | PRJNA688305 | WGS | SRR13318667 |
| ERH95 | MCZ70292 | 44.81 | -117.66 | PRJNA688305 | WGS | SRR13318700 |
| ERH97 | MCZ70294 | 44.81 | -117.60 | PRJNA688305 | WGS | SRR13318678 |
| ERH100 | MCZ70297 | 44.81 | -117.60 | PRJNA688305 | WGS | SRR13318615 |
| ERH110 | MCZ70302 | 44.81 | -117.66 | PRJNA688305 | WGS | SRR13318725 |
| ERH83 | MCZ70286 | 44.85 | -117.64 | PRJNA688305 | WGS | SRR13318714 |
| ERH88 | MCZ70288 | 44.82 | -117.63 | PRJNA688305 | WGS | SRR13318642 |
| ERH89 | MCZ70289 | 44.82 | -117.63 | PRJNA688305 | WGS | SRR13318631 |
| ERH90 | MCZ70290 | 44.85 | -117.64 | PRJNA688305 | WGS | SRR13318758 |
| ERH96 | MCZ70293 | 44.81 | -117.60 | PRJNA688305 | WGS | SRR13318689 |
| ERH107 | MCZ70300 | 44.82 | -117.63 | PRJNA688305 | WGS | SRR13318604 |
| ERH93 | MCZ70291 | 44.81 | -117.66 | PRJNA688305 | WGS | SRR13318747 |
| ERH94 | MCZ69534 | 44.85 | -117.64 | PRJNA688305 | WGS | SRR13318736 |
| **Population f** |  |  |  |  |  |  |
| MSB280706 | MSB280706 | 33.40 | -108.59 | MSB | WGS |  |
| MSB280708 | MSB280708 | 33.40 | -108.59 | MSB | WGS |  |
| MSB280720 | MSB280720 | 33.40 | -108.58 | MSB | WGS |  |
| MSB280729 | MSB280729 | 33.40 | -108.58 | MSB | WGS |  |
| MSB280948 | MSB280948 | 33.40 | -108.58 | MSB | WGS |  |
| MSB280961 | MSB280961 | 33.40 | -108.58 | MSB | WGS |  |
| MSB280963 | MSB280963 | 33.40 | -108.58 | MSB | WGS |  |
| MSB280964 | MSB280964 | 33.40 | -108.58 | MSB | WGS |  |
| MSB280968 | MSB280968 | 33.40 | -108.59 | MSB | WGS |  |
| MSB280977 | MSB280977 | 33.40 | -108.58 | MSB | WGS |  |
| MSB280721 | MSB280721 | 33.40 | -108.58 | MSB | WGS |  |
| MSB280893 | MSB280893 | 33.40 | -108.58 | MSB | WGS |  |
| MSB280973 | MSB280973 | 33.40 | -108.58 | MSB | WGS |  |
| **Population g** |  |  |  |  |  |  |
| SRR8782108 |  | 39.59 | -105.64 | PRJNA528923 | exome | SRR8782108 |
| SRR8782109 |  | 39.59 | -105.64 | PRJNA528923 | exome | SRR8782109 |
| SRR8782110 |  | 39.59 | -105.64 | PRJNA528923 | exome | SRR8782110 |
| SRR8782111 |  | 39.59 | -105.64 | PRJNA528923 | exome | SRR8782111 |
| SRR8782112 |  | 39.59 | -105.64 | PRJNA528923 | exome | SRR8782112 |
| SRR8782113 |  | 39.59 | -105.64 | PRJNA528923 | exome | SRR8782113 |
| SRR8782114 |  | 39.59 | -105.64 | PRJNA528923 | exome | SRR8782114 |
| SRR8782115 |  | 39.59 | -105.64 | PRJNA528923 | exome | SRR8782115 |
| SRR8782116 |  | 39.59 | -105.64 | PRJNA528923 | exome | SRR8782116 |
| SRR8782157 |  | 39.59 | -105.64 | PRJNA528923 | exome | SRR8782157 |
| SRR8782158 |  | 39.59 | -105.64 | PRJNA528923 | exome | SRR8782158 |
| SRR8782159 |  | 39.59 | -105.64 | PRJNA528923 | exome | SRR8782159 |
| SRR8782160 |  | 39.59 | -105.64 | PRJNA528923 | exome | SRR8782160 |
| SRR8782161 |  | 39.59 | -105.64 | PRJNA528923 | exome | SRR8782161 |
| SRR8782162 |  | 39.59 | -105.64 | PRJNA528923 | exome | SRR8782162 |
| SRR8782165 |  | 39.59 | -105.64 | PRJNA528923 | exome | SRR8782165 |
| SRR8782166 |  | 39.59 | -105.64 | PRJNA528923 | exome | SRR8782166 |
| SRR8782167 |  | 39.59 | -105.64 | PRJNA528923 | exome | SRR8782167 |
| SRR8782169 |  | 39.59 | -105.64 | PRJNA528923 | exome | SRR8782169 |
| SRR8782170 |  | 39.59 | -105.64 | PRJNA528923 | exome | SRR8782170 |
| SRR8782171 |  | 39.59 | -105.64 | PRJNA528923 | exome | SRR8782171 |
| SRR8782172 |  | 39.59 | -105.64 | PRJNA528923 | exome | SRR8782172 |
| SRR8782173 |  | 39.59 | -105.64 | PRJNA528923 | exome | SRR8782173 |
| SRR8782174 |  | 39.59 | -105.64 | PRJNA528923 | exome | SRR8782174 |
| SRR8782175 |  | 39.59 | -105.64 | PRJNA528923 | exome | SRR8782175 |
| SRR8782176 |  | 39.59 | -105.64 | PRJNA528923 | exome | SRR8782176 |
| SRR8782177 |  | 39.59 | -105.64 | PRJNA528923 | exome | SRR8782177 |
| SRR8782178 |  | 39.59 | -105.64 | PRJNA528923 | exome | SRR8782178 |
| SRR8782179 |  | 39.59 | -105.64 | PRJNA528923 | exome | SRR8782179 |
| SRR8782180 |  | 39.59 | -105.64 | PRJNA528923 | exome | SRR8782180 |
| SRR8782181 |  | 39.59 | -105.64 | PRJNA528923 | exome | SRR8782181 |
| SRR8782182 |  | 39.59 | -105.64 | PRJNA528923 | exome | SRR8782182 |
| SRR8782183 |  | 39.59 | -105.64 | PRJNA528923 | exome | SRR8782183 |
| SRR8782184 |  | 39.59 | -105.64 | PRJNA528923 | exome | SRR8782184 |
| SRR8782185 |  | 39.59 | -105.64 | PRJNA528923 | exome | SRR8782185 |
| SRR8782186 |  | 39.59 | -105.64 | PRJNA528923 | exome | SRR8782186 |
| SRR8782187 |  | 39.59 | -105.64 | PRJNA528923 | exome | SRR8782187 |
| SRR8782188 |  | 39.59 | -105.64 | PRJNA528923 | exome | SRR8782188 |
| SRR8782189 |  | 39.59 | -105.64 | PRJNA528923 | exome | SRR8782189 |
| SRR8782190 |  | 39.59 | -105.64 | PRJNA528923 | exome | SRR8782190 |
| SRR8782193 |  | 39.59 | -105.64 | PRJNA528923 | exome | SRR8782193 |
| SRR8782194 |  | 39.59 | -105.64 | PRJNA528923 | exome | SRR8782194 |
| SRR8782199 |  | 39.59 | -105.64 | PRJNA528923 | exome | SRR8782199 |
| SRR8782203 |  | 39.59 | -105.64 | PRJNA528923 | exome | SRR8782203 |
| SRR8782204 |  | 39.59 | -105.64 | PRJNA528923 | exome | SRR8782204 |
| SRR8782206 |  | 39.59 | -105.64 | PRJNA528923 | exome | SRR8782206 |
| **Population h** |  |  |  |  |  |  |
| SRR8782117 |  | 40.81 | -96.68 | PRJNA528923 | exome | SRR8782117 |
| SRR8782122 |  | 40.81 | -96.68 | PRJNA528923 | exome | SRR8782122 |
| SRR8782124 |  | 40.81 | -96.68 | PRJNA528923 | exome | SRR8782124 |
| SRR8782125 |  | 40.81 | -96.68 | PRJNA528923 | exome | SRR8782125 |
| SRR8782126 |  | 40.81 | -96.68 | PRJNA528923 | exome | SRR8782126 |
| SRR8782127 |  | 40.81 | -96.68 | PRJNA528923 | exome | SRR8782127 |
| SRR8782128 |  | 40.81 | -96.68 | PRJNA528923 | exome | SRR8782128 |
| SRR8782129 |  | 40.81 | -96.68 | PRJNA528923 | exome | SRR8782129 |
| SRR8782130 |  | 40.81 | -96.68 | PRJNA528923 | exome | SRR8782130 |
| SRR8782131 |  | 40.81 | -96.68 | PRJNA528923 | exome | SRR8782131 |
| SRR8782132 |  | 40.81 | -96.68 | PRJNA528923 | exome | SRR8782132 |
| SRR8782133 |  | 40.81 | -96.68 | PRJNA528923 | exome | SRR8782133 |
| SRR8782134 |  | 40.81 | -96.68 | PRJNA528923 | exome | SRR8782134 |
| SRR8782135 |  | 40.81 | -96.68 | PRJNA528923 | exome | SRR8782135 |
| SRR8782136 |  | 40.81 | -96.68 | PRJNA528923 | exome | SRR8782136 |
| SRR8782137 |  | 40.81 | -96.68 | PRJNA528923 | exome | SRR8782137 |
| SRR8782139 |  | 40.81 | -96.68 | PRJNA528923 | exome | SRR8782139 |
| SRR8782141 |  | 40.81 | -96.68 | PRJNA528923 | exome | SRR8782141 |
| SRR8782142 |  | 40.81 | -96.68 | PRJNA528923 | exome | SRR8782142 |
| SRR8782143 |  | 40.81 | -96.68 | PRJNA528923 | exome | SRR8782143 |
| SRR8782144 |  | 40.81 | -96.68 | PRJNA528923 | exome | SRR8782144 |
| SRR8782145 |  | 40.81 | -96.68 | PRJNA528923 | exome | SRR8782145 |
| SRR8782146 |  | 40.81 | -96.68 | PRJNA528923 | exome | SRR8782146 |
| SRR8782147 |  | 40.81 | -96.68 | PRJNA528923 | exome | SRR8782147 |
| SRR8782148 |  | 40.81 | -96.68 | PRJNA528923 | exome | SRR8782148 |
| SRR8782149 |  | 40.81 | -96.68 | PRJNA528923 | exome | SRR8782149 |
| SRR8782150 |  | 40.81 | -96.68 | PRJNA528923 | exome | SRR8782150 |
| SRR8782151 |  | 40.81 | -96.68 | PRJNA528923 | exome | SRR8782151 |
| SRR8782152 |  | 40.81 | -96.68 | PRJNA528923 | exome | SRR8782152 |
| SRR8782153 |  | 40.81 | -96.68 | PRJNA528923 | exome | SRR8782153 |
| SRR8782154 |  | 40.81 | -96.68 | PRJNA528923 | exome | SRR8782154 |
| SRR8782155 |  | 40.81 | -96.68 | PRJNA528923 | exome | SRR8782155 |
| SRR8782156 |  | 40.81 | -96.68 | PRJNA528923 | exome | SRR8782156 |
| SRR8782163 |  | 40.81 | -96.68 | PRJNA528923 | exome | SRR8782163 |
| SRR8782164 |  | 40.81 | -96.68 | PRJNA528923 | exome | SRR8782164 |
| **Population i** |  |  |  |  |  |  |
| JJK_4190 | MCZ70690 | 38.30 | -84.51 | MCZ | WGS |  |
| JJK_4191 | MCZ70691 | 38.30 | -84.51 | MCZ | WGS |  |
| JJK_4192 | MCZ70692 | 38.30 | -84.51 | MCZ | WGS |  |
| **Population j (*P. m. bairdii*)** |  |  |  |  |  |  |
| MZ10760 |  | 42.20 | -83.89 | Cody Thompson (UM) | WGS |  |
| MZ10762 |  | 42.20 | -83.89 | Cody Thompson (UM) | WGS |  |
| MZ10764 |  | 42.20 | -83.89 | Cody Thompson (UM) | WGS |  |
| MZ10770 |  | 42.20 | -83.89 | Cody Thompson (UM) | WGS |  |
| MZ10772 |  | 42.20 | -83.89 | Cody Thompson (UM) | WGS |  |
| MZ10774 |  | 42.20 | -83.89 | Cody Thompson (UM) | WGS |  |
| MZ10804 |  | 42.20 | -83.89 | Cody Thompson (UM) | WGS |  |
| MZ10808 |  | 42.20 | -83.89 | Cody Thompson (UM) | WGS |  |
| MZ10812 |  | 42.20 | -83.89 | Cody Thompson (UM) | WGS |  |
| MZ10813 |  | 42.20 | -83.89 | Cody Thompson (UM) | WGS |  |
| MZ10814 |  | 42.20 | -83.89 | Cody Thompson (UM) | WGS |  |
| MZ10815 |  | 42.20 | -83.89 | Cody Thompson (UM) | WGS |  |
| MZ10884 |  | 42.20 | -83.89 | Cody Thompson (UM) | WGS |  |
| MZ10885 |  | 42.20 | -83.89 | Cody Thompson (UM) | WGS |  |
| MZ11010 |  | 42.20 | -83.89 | Cody Thompson (UM) | WGS |  |
| MZ11011 |  | 42.20 | -83.89 | Cody Thompson (UM) | WGS |  |
| MZ11012 |  | 42.20 | -83.89 | Cody Thompson (UM) | WGS |  |
| **Population k** |  |  |  |  |  |  |
| EPK001 |  | 45.41 | -84.16 | internal | WGS |  |
| EPK002 |  | 45.41 | -84.16 | internal | WGS |  |
| EPK003 |  | 45.41 | -84.16 | internal | WGS |  |
| EPK004 |  | 45.41 | -84.16 | internal | WGS |  |
| EPK005 |  | 45.41 | -84.16 | internal | WGS |  |
| EPK006 |  | 45.41 | -84.16 | internal | WGS |  |
| EPK007 |  | 45.41 | -84.16 | internal | WGS |  |
| EPK008 |  | 45.41 | -84.16 | internal | WGS |  |
| EPK010 |  | 45.41 | -84.16 | internal | WGS |  |
| EPK011 |  | 45.41 | -84.16 | internal | WGS |  |
| EPK012 |  | 45.41 | -84.16 | internal | WGS |  |
| EPK014 |  | 45.41 | -84.16 | internal | WGS |  |
| EPK017 |  | 45.41 | -84.16 | internal | WGS |  |
| EPK018 |  | 45.41 | -84.16 | internal | WGS |  |
| EPK019 |  | 45.41 | -84.16 | internal | WGS |  |
| **Population l (*P. m. nubiterrae*)** |  |  |  |  |  |  |
| EPK04 |  | 40.33 | -79.27 | internal | WGS |  |
| EPK14 |  | 40.33 | -79.27 | internal | WGS |  |
| EPK07 |  | 40.33 | -79.27 | internal | WGS |  |
| EPK11 |  | 40.33 | -79.27 | internal | WGS |  |
| EPK17 |  | 40.33 | -79.27 | internal | WGS |  |
| EPK08 |  | 40.33 | -79.27 | internal | WGS |  |
| EPK02 |  | 40.33 | -79.27 | internal | WGS |  |
| EPK03 |  | 40.33 | -79.27 | internal | WGS |  |
| EPK01 |  | 40.33 | -79.27 | internal | WGS |  |
| EPK15 |  | 40.33 | -79.27 | internal | WGS |  |
| EPK10 |  | 40.33 | -79.27 | internal | WGS |  |
| EPK18 |  | 40.33 | -79.27 | internal | WGS |  |
| EPK09 |  | 40.33 | -79.27 | internal | WGS |  |
| EPK16 |  | 40.33 | -79.27 | internal | WGS |  |
| NUB13 |  | 40.33 | -79.27 | internal | WGS |  |
| **Population m (*P. p. subgriseus*)** |  |  |  |  |  |  |
| 68144 | MCZ68144 | 29.18 | -81.80 | MCZ | WGS |  |
| 68146 | MCZ68146 | 29.18 | -81.80 | MCZ | WGS |  |
| 68147 | MCZ68147 | 29.18 | -81.80 | MCZ | WGS |  |
| 68148 | MCZ68148 | 29.18 | -81.80 | MCZ | WGS |  |
| 68149 | MCZ68149 | 29.18 | -81.80 | MCZ | WGS |  |
| 68150 | MCZ68150 | 29.18 | -81.80 | MCZ | WGS |  |
| 68151 | MCZ68151 | 29.18 | -81.80 | MCZ | WGS |  |
| 68152 | MCZ68152 | 29.18 | -81.80 | MCZ | WGS |  |
| 68153 | MCZ68153 | 29.18 | -81.80 | MCZ | WGS |  |
| 68154 | MCZ68154 | 29.18 | -81.80 | MCZ | WGS |  |
| 68159 | MCZ68159 | 29.18 | -81.80 | MCZ | WGS |  |
| 68160 | MCZ68160 | 29.18 | -81.80 | MCZ | WGS |  |
| 68161 | MCZ68161 | 29.18 | -81.80 | MCZ | WGS |  |
| 68162 | MCZ68162 | 29.18 | -81.80 | MCZ | WGS |  |
| 68163 | MCZ68163 | 29.18 | -81.80 | MCZ | WGS |  |
| ***P. leucopus*** |  |  |  |  |  |  |
| MSB_212419 | MSB212419 | 36.10 | -94.00 | MSB | WGS |  |
| MSB_212420 | MSB212420 | 36.10 | -94.00 | MSB | WGS |  |
| MSB_212421 | MSB212421 | 36.10 | -94.00 | MSB | WGS |  |
| MSB_53354 | MSB53354 | 46.90 | -96.50 | MSB | WGS |  |
| MSB_53355 | MSB53355 | 46.90 | -96.50 | MSB | WGS |  |
| MSB_53357 | MSB53357 | 46.90 | -96.50 | MSB | WGS |  |
| MSB_229927 | MSB229927 | 43.60 | -73.70 | MSB | WGS |  |
| MSB_229967 | MSB229967 | 43.60 | -73.70 | MSB | WGS |  |
| MSB_42375 | MSB42375 | 33.85 | -100.79 | MSB | WGS |  |
| MSB_42378 | MSB42378 | 33.85 | -100.79 | MSB | WGS |  |
| MSB_42402 | MSB42402 | 33.85 | -100.79 | MSB | WGS |  |
| ***P. californicus*** |  |  |  |  |  |  |
| 47_IS_F_17437 |  | NA | NA | internal | WGS |  |
| IS_SRA |  | NA | NA | PRJNA53595 | WGS | SAMN01081713 |

## Notes

### Competing Interest Statement

The authors have declared no competing interest.

